# AlphaFold-driven discovery of ORP-PIP interactions using new generation confidence scores

**DOI:** 10.1101/2025.09.09.675126

**Authors:** Filippo Dall’Armellina, Sylvie Urbé, Daniel J. Rigden

## Abstract

Non-vesicular lipid transport contributes to the regulation of membrane composition and organelle function at membrane contact sites. OSBP-related proteins (ORPs) are central to this process, yet their interaction networks remain incompletely defined. Here, we systematically screened potential interactions between ORPs and phosphoinositide 3-, 4-, and 5-phosphatases (PIPs) using AlphaPulldown2, AlphaFold2-Multimer, and AlphaFold3. We established a pipeline for model generation by combining AlphaFold2-Multimer predictions (including five-replicates) with an AlphaPulldown2 interaction screen across around 200 protein pairs, and with AlphaFold3 predictions including lipid-bound and multimeric assemblies. Interface confidence was assessed for consistency using the weighted ipTM+pTM metric, actifpTM, new generation ipSAE scoring, and FoldSeek-Multimer clustering. We further evaluated the protein pairs’ biological plausibility based on subcellular localisation data, *in silico* membrane insertion, evolutionary conservation via ConSurf, and protein binding interface analysis using the deep learning tool PeSTo. This integrative pipeline uncovered functionally conserved binding modes in the SAC1 lipid phosphatase with the ORP family, particularly with ORP11, and predicted functionally relevant protein-lipid interfaces.

## 1 Introduction

Membrane contact sites (MCSs) are specialised subcellular compartments where organelles are closely apposed, typically 10-40 nm apart, enabling non-vesicular exchange of lipids, ions, and metabolites ^1,2^. Transfer between membranes can occur through protein-mediated mechanisms (shuttle or bridge models), vesicular transport, or, when membranes approach within 2-3 nm, direct lipid diffusion ^1,3^. Cargo includes ions, metabolites, and lipids. Among the lipid cargo, phosphoinositides (PIs) are particularly important. These phospholipids act both as precursors for second messengers and as organisers of protein complexes on organelle membranes ^4,5^. PIs regulate processes including membrane architecture, cytoskeletal dynamics, and signal transduction ^4,6,7^. Their functions depend on both localisation and phosphorylation state. Kinases and phosphatases act at positions 3, 4, and 5 of the inositol ring to generate seven phosphorylated species ^8^.

Phosphatases are widely distributed across organelles, but several function at the endoplasmic reticulum (ER) or plasma membrane (PM), where many MCSs form. Well-studied examples include: (1) OCRL, a 5-phosphatase that functions within the secretory pathway, localising to the trans-Golgi network and, under certain signals, to the PM ^9^; (2) INPP5K, another 5-phosphatase, recruited to the ER surface by ARL6IP1 to hydrolyse PI(4,5)P2 ^10^ and also detected at the PM in signalling contexts ^11^; and (3) SAC1, an ER-embedded 4-phosphatase that hydrolyses PI4P molecules ^12^. Together, these enzymes illustrate how phosphatases not only remove phosphate groups but also actively shape the lipid environment, preparing lipids for transfer at MCSs ^3^.

The ER-PM interface is a key MCS hosting many lipid transfer proteins ^3^, including the oxysterol-binding protein (OSBP)-related proteins (ORPs) which mediate non-vesicular lipid exchange between organelles ^13-17^. ORPs are typically recruited to MCSs through interactions with VAMP-associated proteins (VAPs) at the ER or with phosphatidylinositol 4-phosphate (PI4P)-enriched membranes ^16^, such as the PM or Golgi apparatus. Most ORPs contain a canonical FFAT motif (two phenylalanines in an acidic tract) that binds to the major sperm protein (MSP) domain of VAPA/B transmembrane (TM) proteins embedded in the ER ^14,18,19^. Beyond this, ORPs share three main domains: (1) a coiled-coil domain, enabling dimerisation (e.g., ORP9– ORP11) ^13,17^; (2) a pleckstrin homology (PH) domain; and (3) an OSBP-related lipid-binding (ORD) domain, accommodating ligands including sterols and Pis ^15,20^.

ORPs can exchange cholesterol derivatives for other lipids across membranes and often functionally cooperate with SAC1, which dephosphorylates PI4P at ER-PM contact sites ^21^. Together, ORPs and SAC1 generate and maintain lipid gradients across intracellular compartments ^21,22^. SAC1 was initially proposed to act *in trans* ^23^, but subsequent studies suggest it acts *in cis* instead ^24,25^. *In cis* means that SAC1 dephosphorylates PI4P after the lipid has been transferred into the same membrane SAC1 is embedded in, the ER. Thus, PI4P is first extracted from the plasma membrane (PM) by an ORP, transferred to the ER, and then dephosphorylated by SAC1 within the ER membrane. *In trans*, SAC1 would act directly on PI4P located in the opposing membrane, the PM, without the lipid being transferred first. This requires SAC1’s catalytic site to reach the adjacent membrane at an ER-PM contact site. Given that SAC1 is an ER-resident membrane protein, the “cis”-model is structurally and biologically more plausible ^23,24^.

Given the central role of PI metabolism at MCSs, we hypothesised that transient or cooperative interactions between PI 3-, 4-, and 5-phosphatases (PIPs) and lipid transfer proteins may facilitate substrate handoff or spatial regulation during cellular signalling events. We designed a modelling pipeline with two state-of-the-art structural prediction tools: AlphaPulldown2 and AlphaFold3. AlphaPulldown2, built on AlphaFold2-Multimer, offers a scalable framework for all-against-all interaction screening ^26^. We treat AlphaFold2-Multimer and AlphaFold3 as completely separate tools. AlphaFold2-Multimer was used because it integrates scoring metrics beyond the default ipTM score: ipSAE and actifpTM ^27,28^. By integrating outputs from both *in silico* methods and evaluating models using multiple confidence metrics (pLDDT, PAE, ipSAE, actifpTM), we mapped a landscape of potential interactions and contextualised them with subcellular localisation datasets.

## 2 Methods

### 2.1 Protein structure prediction

Structural predictions were performed using a local installation of LocalColabFold ^29,30^ on a high-performance workstation running Ubuntu 24.04.1 LTS (kernel 6.8.0-51-generic). The machine was equipped with an AMD Ryzen Threadripper 3970X 32-core processor (64 threads), 125 GiB RAM, and an NVIDIA GeForce RTX 3080 GPU with 10 GB VRAM, managed via NVIDIA-SMI 535.183.01 and CUDA 12.2. All computations were executed in a Conda environment running Python 3.9 and the necessary dependencies. GPU usage during AlphaFold2 runs was modest, with approximately 588 MiB of VRAM used, mostly by system processes.

AlphaFold2 outputs several diagnostic plots and confidence metrics to assess the quality of its structural predictions. One such plot displays the coverage and depth of the initial multiple sequence alignment (MSA), alongside a map of regions covered in the full-length protein. Among the key confidence scores is the predicted Local Distance Difference Test (pLDDT), which provides a per-residue estimate of model accuracy on a scale from 0 to 100. Scores above 90 indicate very high local accuracy, comparable to experimentally determined structures, allowing reliable interpretation of side-chain positioning. Scores between 70 and 90 reflect confident backbone predictions, while values between 50 and 70 suggest moderate confidence, where overall secondary structure elements (e.g., α-helices and β-strands) are likely correct, but their relative arrangement may be uncertain.

In addition to pLDDT, AlphaFold2 generates a Predicted Aligned Error (PAE) plot, which estimates the positional error (in Angstroms) between all residue pairs in the model. The PAE is visualised as a 2D heatmap, where lower values indicate high confidence in the spatial relationship between residues. Typically, PAE values are low in well-folded domains and higher between domains that may be flexibly linked or lack defined contact, reflecting uncertainty in their relative orientation. This makes PAE particularly useful for identifying flexible linkers or domain-domain arrangements that may be functionally important but structurally dynamic.

AlphaFold2 predictions in this study incorporated a refined confidence metric known as actual interface predicted TM-score (actifpTM) ^28^. This enhancement addresses a key limitation of the original interchain predicted TM-score (ipTM), which struggles to accurately assess interaction interfaces in the presence of long intrinsically disordered regions or non-interacting flanking segments, features commonly omitted in crystal structures but often present in full-length sequences. While ipTM has been applied in docking studies involving peptide-protein complexes ^31-33^, it was originally trained on structured PDB entries and is sensitive to sequence length, treating all residues equally regardless of their involvement in the interface. As a result, inclusion of disordered or accessory regions can lead to substantial reductions in ipTM scores, even when the core interaction is well predicted.

To mitigate this, actifpTM modifies the ipTM calculation by masking non-interacting regions and reweighting residue pairs based on predicted distance probabilities. This makes the score more reflective of true interface quality, independent of overall sequence length or structural disorder. Two components are essential for calculating the actifpTM score, both related to contact maps derived from distograms generated during model prediction. Note that a distogram is required to calculate the contact map. AlphaFold3 does not provide distograms, so it cannot calculate this score from a model. Second, probability maps from the PAE matrix are needed. These are not produced in the AlphaFold3 pipeline ^34^.

The ipTM evaluates interactions between chains rather than individual interface regions. To address this, we employed the ipSAE ^27^. The ipsae.py script calculates ipSAE based entirely on interchain residue pairs with PAE(i,j) values below a defined cutoff. When no residue pairs meet this criterion, ipSAE returns 0. The script also computes two additional forms for comparison: ipSAE_d0chn and ipSAE_d0dom. ipSAE_d0chn uses the same PAE cutoff as ipSAE but defines d0 as the sum of the full-length sequences of the two chains. ipSAE_d0dom calculates d0 from the number of residues in the two chains that have any interchain PAE values below the cutoff. Importantly, ipSAE does not require modifications to the AlphaFold codebase and relies solely on standard output files (e.g., JSON) from both AlphaFold2-Multimer and AlphaFold3. A PAE cutoff of 10 Å and an ipSAE threshold of 0.3 have been suggested to distinguish likely true interactions from false positives ^27^. For comparisons across ORP-PIP complexes, we focus on ipSAE_d0dom rather than the standard ipSAE score because it normalises by residues that participate in the interchain interface. This prevents differences in chain length or disordered regions in the ORP and PIP sequences from artificially lowering the confidence score, enabling a more accurate comparison of potentially conserved interfaces across structurally aligned models, which are subsequently analysed with FoldSeek-Multimer ^35^.

For completeness, we also used the well-established pDockQ score. pDockQ, built for AlphaFold2-Multimer outputs, is a *predicted* DockQ score that does not require a reference structure ^36^. Instead, it estimates DockQ directly from model-internal metrics, including predicted IDDT and interface properties. The resulting continuous score, ranging from 0 (low quality) to 1 (high quality), provides a proxy to the reference DockQ ^37^ and was used here to give a comprehensive overview of the models by running the ipSAE script ^27^.

For AlphaFold3 predictions, we used the open-access AlphaFold Server released in mid-2024 ^34^, which enables modelling of complex multimolecular assemblies, including proteins, peptides, and nucleic acids. Structural models were examined using the AlphaFold3 Visualiser script ^38^ to inspect residue-level interactions. For multimeric predictions, we further analysed inter-chain contacts using PDBePISA ^39,40^. All models were visualised and interpreted in PyMOL (version 2.1.1, Schrödinger, LLC, New York, NY, USA), which was used for figure generation and manual inspection of predicted interfaces. The ORP11-SAC1 complex was further analysed for its interface residue data using AlphaBridge ^41^. AlphaBridge is a tool that identifies and characterises residue-residue interactions in protein complex models generated by AlphaFold.

### 2.2 Analysis of AlphaPulldown2 interaction screens

To systematically evaluate potential protein-protein interactions at scale, we employed AlphaPulldown2 ^26^, a high-throughput wrapper for AlphaFold-Multimer designed for all-against-all structural interaction screens. AlphaPulldown2 automates the generation of model inputs, prediction jobs, and post-processing, making it particularly well-suited for matrix-style interaction mapping. The all-by-all screening configuration results in predictions evaluated using a custom confidence pipeline that integrated multiple structural metrics, including pLDDT, PAE, actifpTM ^28^, pDockQ ^36,42^, ipSAE ^27^ and FoldSeek-based multimer clustering to detect recurring interface patterns across models ^35,43,44^.

We generated complexes for nearly 200 protein pairs using AlphaPulldown2, with five replicates per pair. From here, we employed: (1) AlphaFold2-Multimer to calculate actifpTM scores; and (2) AlphaFold3 to refine the top AlphaPulldown hits (ipTM+pTM > 0.5). AlphaFold2-Multimer and AlphaFold3 were also used to model higher-order assemblies (3-4 protein chains), including examples such as dimers bound to a phosphatase protein. Note that protein-lipid complexes were additionally generated with the AlphaFold3 server using palmitic acid (PLM) as a proxy for lipid binding.

### 2.3 Construction of protein-membrane simulation systems

The process of building a membrane bilayer-protein system via CHARMM-GUI involves several steps and was conducted using Bilayer Builder in Membrane Builder ^45-47^. It begins with loading protein coordinates and uploading a pre-oriented protein structure ^48^. This was followed by determining the system size with a heterogeneous rectangular lipid bilayer, with a specified water thickness parameter of 10 Å and a defined ratio of lipid components (no generation of pore water). The replacement method was then used to build the system, omitting the ion-placing method ^49^. System equilibration and relaxation were performed applying to the system the CHARMM36 lipid force field with WYF parameters to accurately capture cation-π interactions between aromatic amino acids and cations and using the automatic PME FFT grid generation method to calculate long-range electrostatic interactions in periodic systems. The system was subjected to a thermodynamic NPT ensemble, maintaining a physiological temperature of 310.15 K.

### 2.4 Functional interface prediction coupled with ConSurf and PeSTo

To assess the potential functional relevance of candidate interfaces, we applied the ConSurf method to identify evolutionarily conserved regions ^50^. This approach allowed us to highlight residues likely to be functionally important or structurally constrained. In parallel, we used PeSTo ^51^, a parameter-free geometric deep learning tool, to predict protein-protein interaction interfaces. By integrating ConSurf conservation data with PeSTo’s interface predictions, we aimed to pinpoint conserved regions likely involved in mediating interactions between the proteins of interest.

### 2.5 Subcellular localisation analysis from proteomics data

We consulted the Human Cell Map non-negative matrix factorisation (NMF) outputs ^52^, Prolocate ^53^ and Map of the Cell ^54^ databases for our proteins of interest. The data we extracted from Prolocate came from Experiment B, conducted on liver tissue of fed rats. When proteins of interest were not detected under these normal conditions, we relied on annotations from Experiment A on fasted rats.

## 3 Results

### 3.1 New generation scores reveal ORP-PIP interactions

We systematically screened 12 human ORPs for potential interactions with 16 PIPs implicated in PI turnover at ER-PM and ER-Golgi junctions. Our goal was to identify potential interactions between PIPs and lipid transfer proteins and explore how structure and localisation data converge to suggest functional partnerships. To this end, we created a pipeline for systematic structural screening (**Fig. 1A**). This effort produced a structural interaction dataset of nearly a thousand AlphaPulldown2 models. The top-scoring structures were identified, and their sequences were independently modelled and rescored using AlphaFold3. Despite substantial variability between model confidence scores across methods, a subset of interactions consistently scored above threshold and showed spatial interface coherence, suggesting genuine biological relevance.

**Fig. 1.**
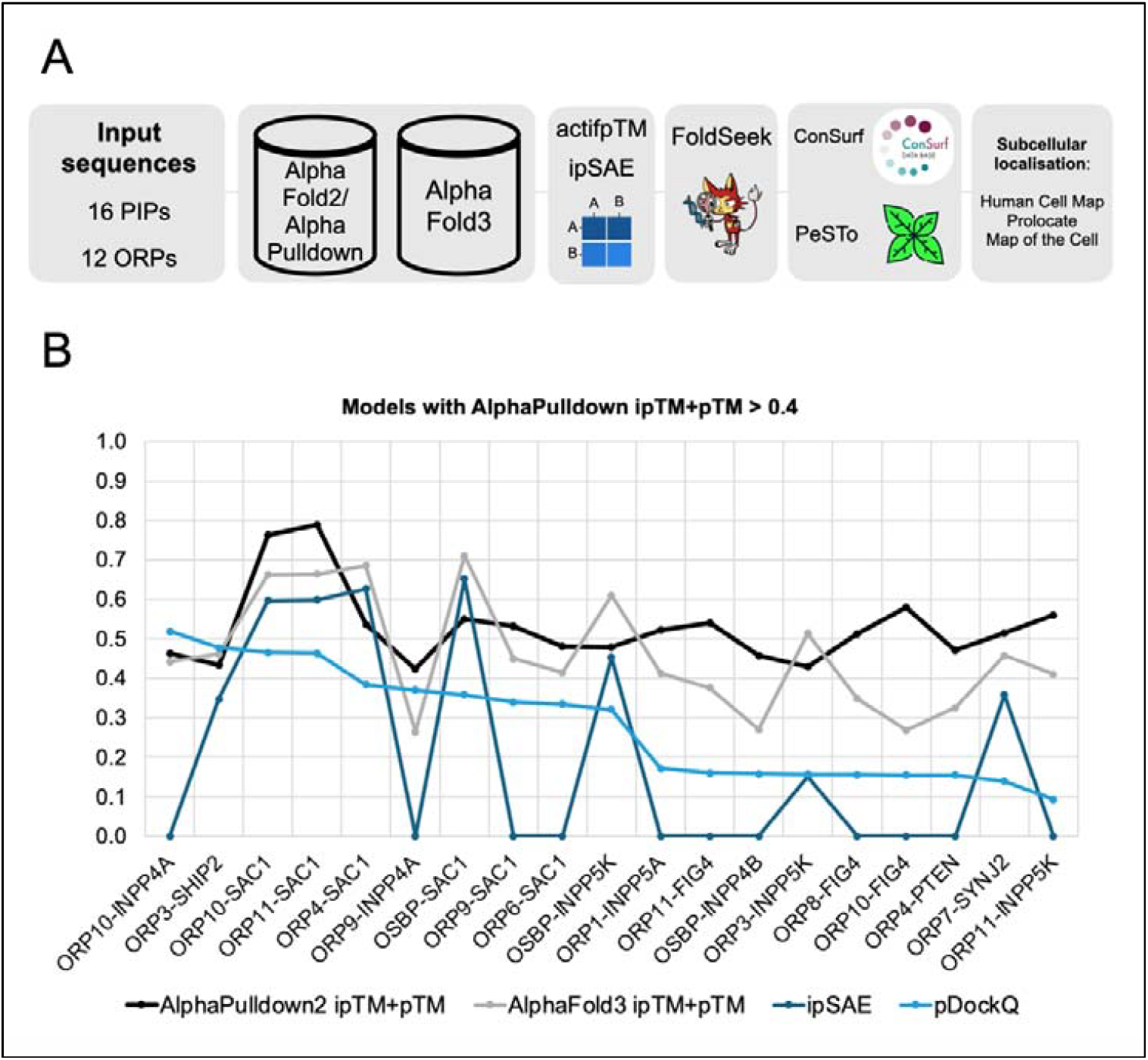
**A.** Structural screening pipeline for ORP-PIP interactions. We employed AlphaFold3 and AlphaFold2-Multimer/AlphaPulldown2, both support ipSAE scoring but only AlphaFold2-Multimer can calculate actifpTM scores. Models were structurally aligned against each other with FoldSeek-Multimer, and then consensus interactions were analysed using ConSurf, PeSTo, and subcellular localisation datasets (i.e. Prolocate, Map of the Cell) to highlight functionally relevant partnerships. “ipTM+pTM” represents the weighted confidence score, calculated as 0.8 × ipTM + 0.2 × pTM. “ipSAE” represents the ipSAE_d0dom score. **B**. Graph of AlphaPulldown and AlphaFold3 models from the ORP dataset. 12 human paralogues were used as baits against 16 PIPs to predict protein-protein interactions.

In an initial AlphaPulldown2 screen we applied a permissive weighted confidence ipTM+pTM threshold (>0.5) to maximise sensitivity for our models of interest. **Fig. 1B** shows a 3D scatter plot where 22 models exceed this cutoff shown on the z-axis. The dot colour encodes the corresponding AlphaFold3 weighted confidence score, revealing a relatively poor correlation between AlphaPulldown2 and AlphaFold3. Several models scored highly in one system but not the other, like in the case of pairs involving the phosphatase INPP4A. These discrepancies highlight the limitations of relying on a single confidence metric.

Those models scoring high with ipSAE also showed high-confidence values in AlphaFold3 (shown in green), suggesting the ipSAE score might correlate better with the more holistic confidence estimation in AlphaFold3 (**Fig. 1B**). Still, we must remain cautious, particularly with the weighted ipTM+pTM metric, which penalises models with low pTM, which can result from disordered regions, present in some of the proteins studied here. This is especially evident in the ORP protein chains, where flexible linkers reduce global confidence despite strong local interfaces.

### 3.2 Interface clustering reveals SAC1 can interact with ORP paralogues

While ORPs vary widely in domain architecture and organelle specificity according to the literature ^14,16,18,19^, many shared a similar orientation when binding to SAC1 in our models. Therefore, we performed systematic clustering of predicted complexes based on interface similarity rather than global structure or sequence identity. This approach uncovered a conserved SAC1-binding mode across multiple ORP paralogues. To examine any inconsistencies between AlphaPulldown2 and AlphaFold3 modelling and scoring, we introduced FoldSeek-Multimer clustering to cross-check interfaces predicted independently by the two tools. Since our top-scoring models involved protein pairs containing INPP5K or SAC1, we focused on this subset and trimmed the ORP models to their ORD domains after observing that this region consistently formed the primary interface across the dataset.

Using the FoldSeek-Multimer cluster pipeline, we calculated interface-local lDDT scores for predicted complexes involving all 12 ORPs and our panel of 16 PI phosphatases. To ensure we captured meaningful structural convergence, we set an arbitrary interface similarity cutoff (interface lDDT > 0.8), similar to that used in global alignments with the TM-score ^55^. At this threshold, only models involving SAC1 retained strong clustering, suggesting a structurally conserved binding interface not seen with predicted complexes involving other phosphatases (**Fig. 2**).

**Fig. 2.**
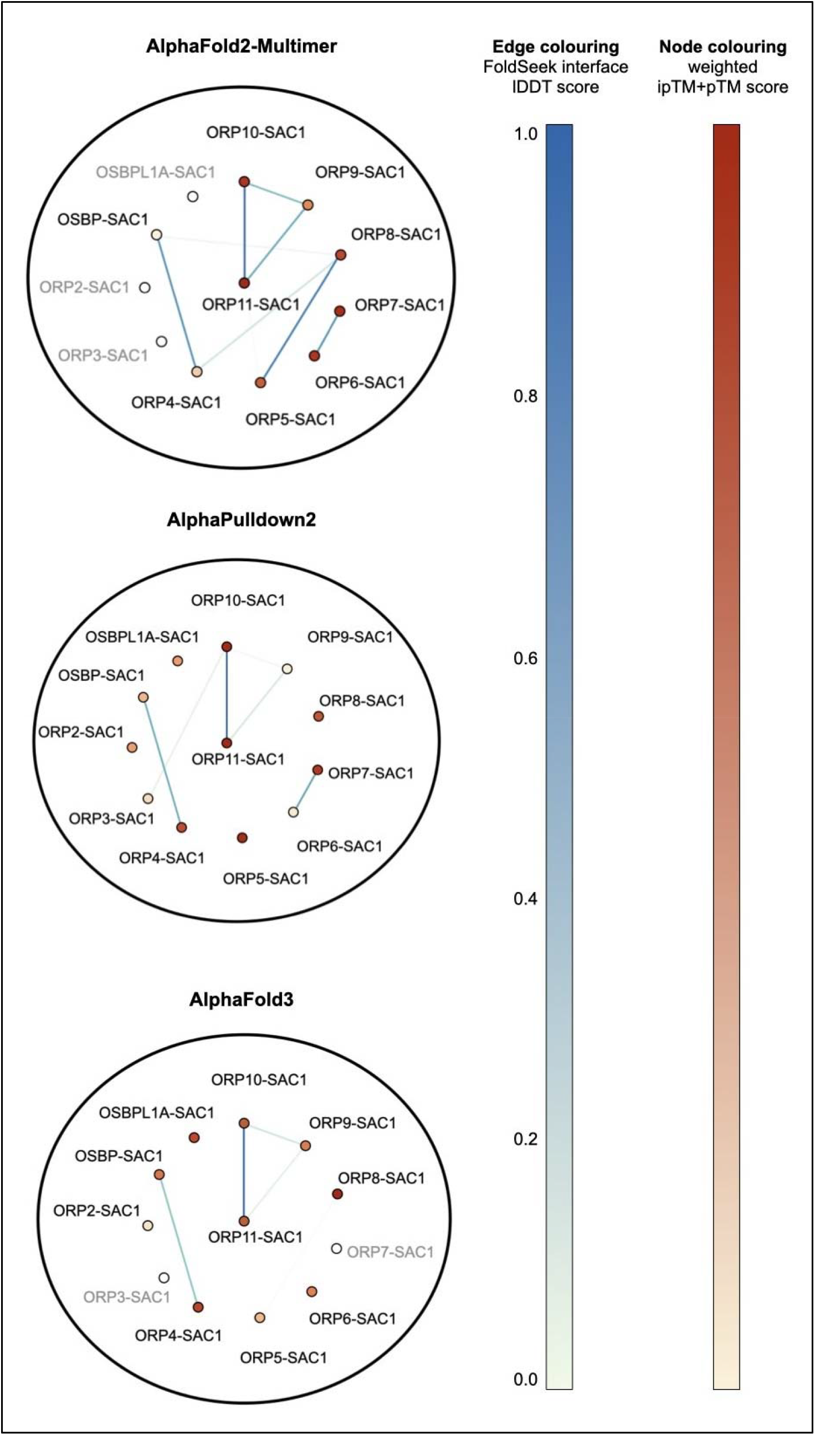
Network clustering of SAC1-ORP protein pairs based on interface similarity using FoldSeek-Multimer. Three networks were generated from models produced by AlphaFold2-Multimer, AlphaPulldown2, and AlphaFold3. Models were trimmed to the ORP ORD domain, while SAC1 was kept full-length. Each node represents a model with ipTM+pTM > 0.5. Edges reflect interface lDDT similarity, with thickness and colour indicating similarity strength and ipTM+pTM scores (blue and orange scales, respectively). Networks were filtered for FoldSeek interface lDDT > 0.8. Grey text boxes represent complexes that were not modelled in AlphaFold2-Multimer or AlphaFold3 as the same protein pairs in AlphaPulldown2 had a weighted ipTM+pTM confidence score < 0.5. Layout: Fruchterman-Reingold algorithm; analysed in Gephi ^56^.

We visualised the similarity network using a Fruchterman-Reingold layout, where each node represents a model and edges reflect interface similarity scores. Models formed a scattered network. If we apply an lDDT>0.8 threshold, a coherent subcluster emerges, consisting of SAC1 bound to ORP10 and ORP11. This refined cluster is consistent across the different versions of AlphaFold we used: the nodes indicate strong ipTM+pTM scores and the edges highlight consistent interface geometries between the SAC domain of SAC1 and the ORD domain of ORPs. Importantly, we took a conservative approach in interpreting the data, focusing only on models where both interface similarity and confidence metrics were significant. A level of agreement between AlphaFold3, AlphaFold2-Multimer, and AlphaPulldown2 models for individual protein pairs, was seen only for the SAC1-ORP11 complex. These results underscore the added value of interface-based metrics in large-scale structural screens.

We further illustrate the discrepancies between AlphaPulldown and AlphaFold3 in modelling the ORP9-SAC1 (**Fig. S1**) and ORP10-SAC1 complexes (**Fig. S2**). The primary differences are in the coiled-coil domain region and the relative orientation of the PH domain, both of which shift substantially between the two modelling approaches, and it is possible to see the structural similarities across protein pairs clearly after removing these regions. These domain-level rearrangements introduce noise into the FoldSeek-Multimer clustering results. **Fig. 2** includes cross-ORP and cross-program comparisons (i.e., AlphaPulldown vs. AlphaFold3), revealing that while some models appear visually similar especially at the interface, their local interface lDDT scores occasionally fall just below our 0.8 threshold. We note that this threshold would have to be much lower if we clustered our protein pairs without trimming the ORP models down to their ORD domains, as FoldSeek was less accurate in aligning structures.

Using our approach, several previously unreported ORP-SAC1 models sharing similar structures emerged, including ORP4-SAC1/OSBP-SAC1, ORP10-SAC1/ORP11-SAC1, and ORP9-SAC1/ORP10-SAC1. These new interactions consistently appeared across all three modelling methods. While we do not analyse each complex in depth, this highlights the value of applying a “trimming” strategy to interface structural alignments as a practical way to leverage FoldSeek-Multimer and expand the landscape of plausible interactions.

We observed strong agreement for ORP6-SAC1 and ORP7-SAC1 models between AlphaPulldown2 and AlphaFold2-Multimer, and for ORP5-SAC1 and ORP8-SAC1 between AlphaFold2-Multimer and AlphaFold3. These predictions, based solely on the ORD domain of ORPs, suggest a conserved binding interface. We focused on ORP11-SAC1 above other pairs, and subsequently on its similarities with ORP9-SAC1 and ORP10-SAC1, since the other AlphaFold models show greater uncertainty in the orientation of PH and coiled-coil domains, whereas ORP11 consistently displays a clear SAC1 binding interface across AlphaFold methods. Notably, the predicted orientation of SAC1’s ER-embedded TM domain in contact with ORP4, ORP6, and ORP7 coiled-coil domains is biologically implausible, and we do not consider it a valid secondary binding site.

### 3.3 ORP11-SAC1 in a membrane system

From our broad screen of ORP-phosphatase interactions, the ORP11-SAC1 complex emerged as a particularly robust pairing, with consistent high-confidence models across AlphaFold2, AlphaFold3, and FoldSeek-Multimer clustering. Building on this, we manually superimposed the AlphaPulldown2 and AlphaFold3 models of ORP11-SAC1, both of which score above 0.8 in interface lDDT according to FoldSeek-Multimer, to assess their structural agreement in more detail.

We noticed that while the core interface between the SAC1 catalytic domain and the ORP11 ORD domain aligns closely across models, notable differences appear elsewhere (**Fig. 3A-C**). The coiled-coil α-helices of ORP11 occupy different positions in each model, and the PH domain undergoes a complete shift in orientation. These discrepancies align with regions of lower predicted alignment confidence in the PAE heatmaps, particularly those corresponding to the sequences of the N-terminal half of ORP11 domains (**Fig. 3A-C**). In contrast, the core SAC1-ORD interface lies in a high-confidence region, reinforcing the consistency of this interaction. Colouring the models by pLDDT shows that AlphaPulldown2 and AlphaFold3 differ in this same region, consistent with the lower predicted alignment confidence in the PAE heatmaps just discussed (**Fig. S3B**). While both AlphaPulldown2 and AlphaFold3 assign high pLDDT scores to the core interface, AlphaPulldown2 exhibits substantially lower confidence in the coiled-coil and PH domains compared to AlphaFold3 (**Fig. S3B**). This suggests that AlphaFold3 may provide a more reliable model overall, at least for these flexible or peripheral regions. Despite these local uncertainties, the main SAC1-ORD interface remains structurally the same between methods, underscoring the robustness of this interaction even when peripheral domain positions diverge. The SAC1-ORD interface comprises three key interactions as reported by AlphaBridge results for the AlphaFold3 model (**Fig. 3D**) and more than 90% of interacting residues of SAC1 and ORP11’s ORD domain are shared between models generated by AlphaFold3, AlphaFold2-Multimer, and AlphaPulldown2.

**Fig. 3.**
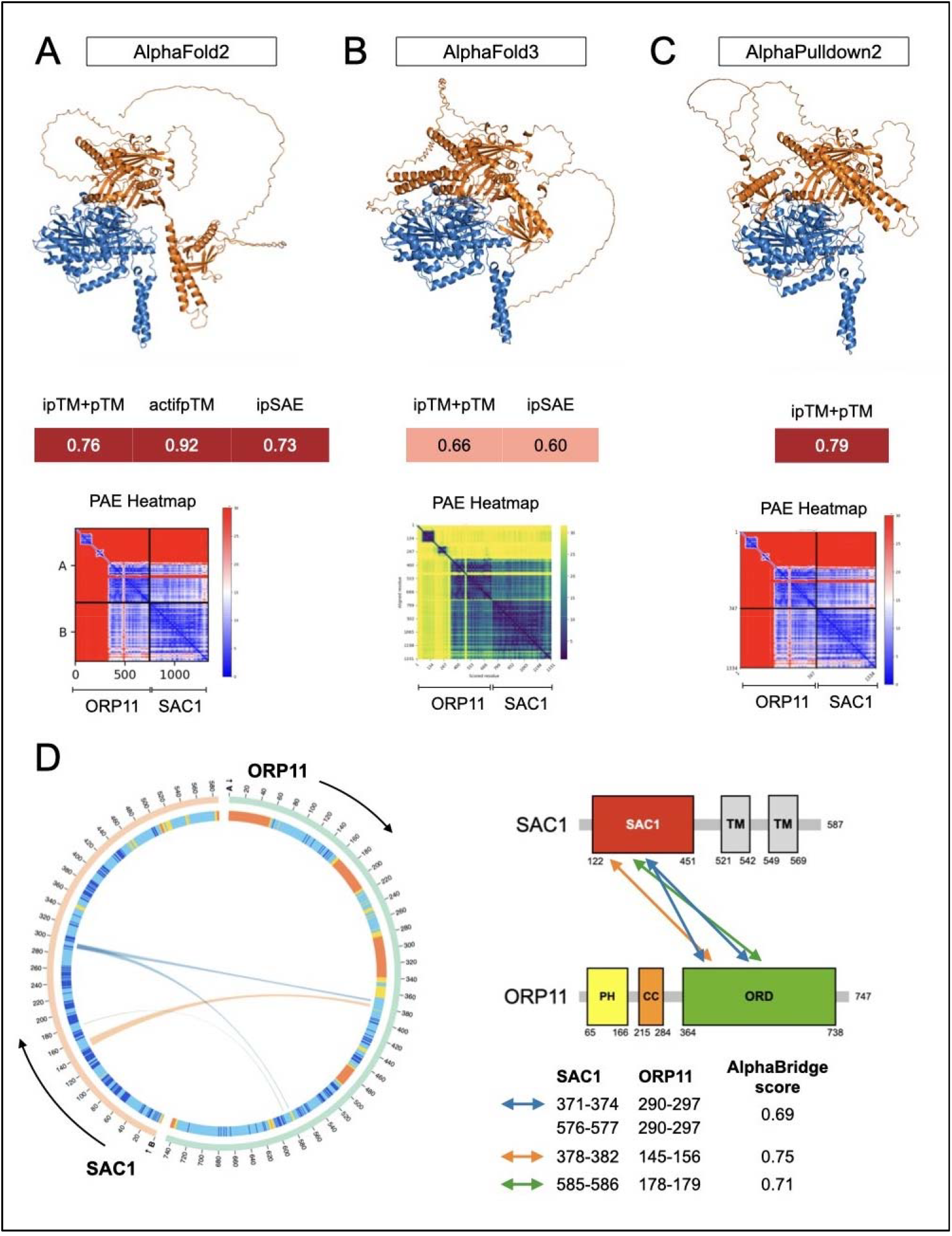
Structural superimposition of **A**. AlphaFold2-Multimer, **B**. AlphaFold3, **C**. and AlphaPulldown2 models for the ORP11-SAC1 complex. PAE heatmaps and confidence scores for each model are shown right below. “ipTM+pTM” represents the weighted confidence score, calculated as 0.8 × ipTM + 0.2 × pTM. “ipSAE” represents the ipSAE_d0dom score. **D**. AlphaBridge results for the AlphaFold3 ORP11-SAC1 model ^41^. Left: chord diagram showing residue-residue interactions between ORP11 and SAC1 along a circular representation of their sequences, coloured by per-residue pLDDT confidence scores (AlphaFold colour scheme; credit: Konstantin Korotkov). Lines connecting residues indicate contacts at the interface and are coloured according to the AlphaBridge interaction score, reflecting interaction strength. Right: protein domain maps of ORP11 and SAC1, with three arrows (blue, orange, and green) highlighting the regions involved in the interaction.

Having established a preference for the AlphaFold3 model of ORP11-SAC1 based on the near-ideal positioning of the PH domain adjacent to the putative membrane plane where SAC1 resides, we next questioned whether the PH domain was oriented appropriately relative to its bound lipid. In **Fig. S3A**, we superimposed the lipid-bound PH domain from ARHGAP9 (PDB: 2P0D) onto the ORP11 PH domain in each model to assess its compatibility with membrane binding. In the AlphaFold3 model, the PH domain is oriented toward the membrane in a configuration compatible with canonical lipid docking. In contrast, the AlphaPulldown2 model positions the PH domain closer to the SAC domain, possibly indicating an alternate, transient or less reliable interface. Despite these differences, both models share most key interface residues that overlap between SAC1 and the ORP11 ORD domain.

We next explored AlphaFold3’s recently discussed capacity to model protein-ligand interactions ^57^, despite its current lipid repertoire being limited to palmitic acid (PLM) and oleic acid. We aimed to assess whether these ligands could approximate the positioning and binding behaviour seen in published co-crystal structures containing PI lipids, particularly in relation to the PH domain, ORD domain, and SAC1. To test this, we generated an ORP11-SAC1 complex model with three PLM molecules: one for each lipid-binding domain. AlphaFold3 did not place a PLM in the PH domain of ORP11 but instead docked two ligands into the lipid-binding groove of the ORD domain, engaging regions that overlap with those observed in experimental co-crystal structures (**Fig. S4B**). Confidence scores for these models were higher than those of the lipid-free AlphaFold3 complex (**Fig. 3B, S4A**). One of the PLM molecules is very closely positioned near the catalytic site of SAC1, highlighted in **Fig. S4A**.

Finally, we systematically increased the number of PLM molecules to explore potential lipid-binding sites, generating models with 1, 2, 3, 5, 10, 15, 25, and finally 50 PLMs, the maximum permitted by the AlphaFold3 server. As the number of PLMs increased, model confidence (ipTM+pTM) declined, largely due to poor ligand placement and the presence of unstructured contacts. In the 50-PLM model, most ligands clustered around SAC1’s TM region, consistent with PeSTo-predicted lipid-binding sites (**Fig. 4D**), but many failed to engage meaningfully with the protein surface, contributing to the reduced confidence scores. Additionally, the PH and coiled-coil domains of ORP11 showed high positional variability across models, further impacting structural confidence (**Fig. S4C**).

**Fig. 4.**
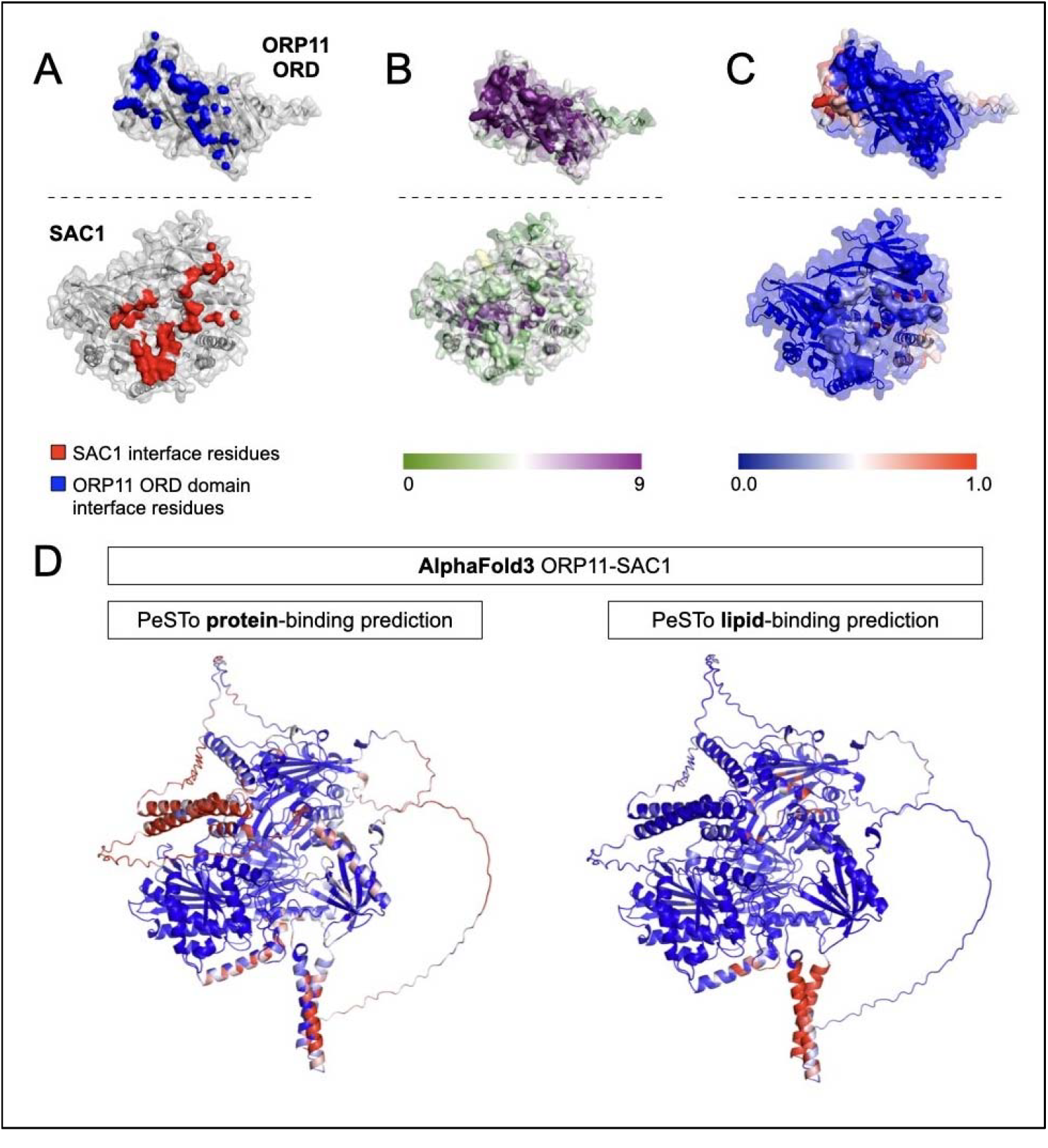
**A.** Open-book view of the ORP11 ORD-SAC1 interface, highlighting key interacting residues. **B**. Same view coloured by evolutionary conservation from ConSurf. **C**. Same view showing PeSTo-predicted protein-protein binding residues. **D**. Full complex view (same orientation as in **Fig. 3**), with PeSTo predictions overlaid: protein-protein binding residues on the left and lipid-binding residues on the right.

When a small number of PLM molecules were included, we observed a moderate improvement in interface confidence scores, including weighted ipTM+pTM and pLDDT, suggesting that the presence of a hydrophobic ligand helps stabilise the predicted complex and promotes the formation of a more realistic lipid-binding cavity. However, increasing the lipid count to 25 or even 50 molecules, led to a sharp drop in overall confidence as indicated by the weighted ipTM+pTM scores, unrealistic crowding in the binding pockets, and diffusion of the model’s attention across too many uncoordinated ligands. We interpret this as AlphaFold3 exhausting plausible, high-confidence binding sites, leading to non-specific or surface-exposed placements, mimicking the bilayer by forming bulk interactions. Score fluctuations emphasise the sensitivity of AlphaFold3 to domain mobility and ligand-induced conformational changes, even when the underlying interface remains largely similar.

To assess the potential functional relevance of the ORP11-SAC1 interface, we visualised the predicted interface using an open-book view (**Fig. 4A**). Conservation analysis with ConSurf (**Fig. 4B**) indicated that several interface residues, particularly on ORP11, are evolutionarily conserved, with some conserved residues present on SAC1. Protein-protein interaction predictions from PeSTo (**Fig. 4C**) supported these findings. Notably, the following residues on ORP11 were identified by both ConSurf and PeSTo as likely interface sites: S368, V369, H372, L373, and Q376-D382. For SAC1, overlapping predictions included Y482 and K483, though overall agreement between the two methods was lower for this protein. There is a high-scoring helix in SAC1 predicted to interact with both proteins and lipids (**Figs. 4D**). The helix runs parallel to the membrane bilayer, and when SAC1 is embedded at the ER, its proximity to the upper leaflet likely occludes any potential protein binding sites.

We used CHARMM-GUI to assess the compatibility of our models with membrane insertion (**Fig. 5A**). Residues within 3.5 Å of the membrane surface are highlighted in green. In the AlphaFold3 model, the PH domain of ORP11 sits just above the membrane, poised for lipid engagement, while the ORD domain contacts SAC1 laterally. In contrast, the AlphaPulldown2 model positions the PH domain away from the membrane, possibly reflecting an alternate or pre-bound state. Nevertheless, in both the AlphaFold3 and AlphaPulldown2 models, the PH domain remains close enough to the membrane for lipid binding without being occluded or hindered by SAC1 (**Fig. 5B**).

**Fig. 5.**
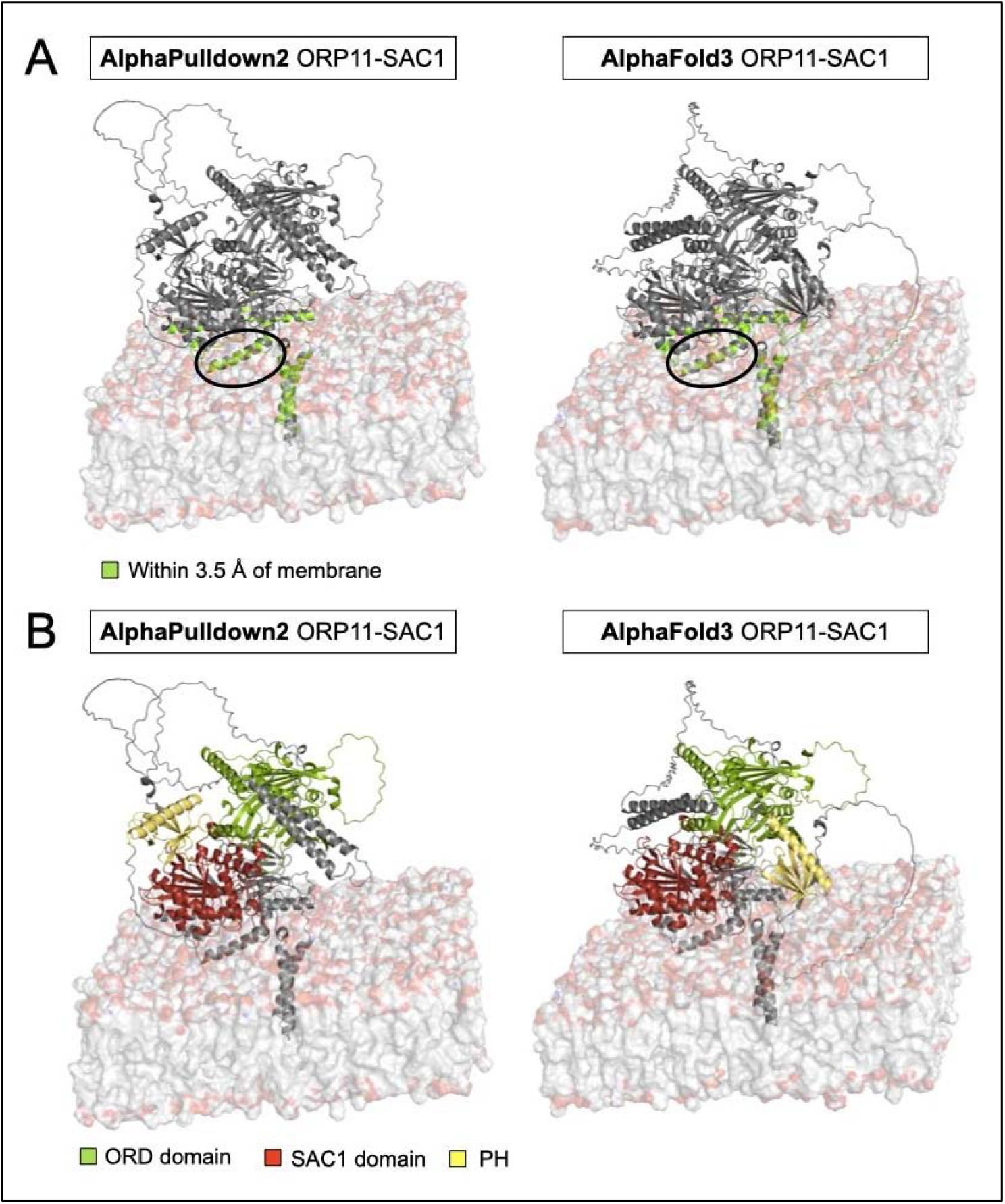
**A.** AlphaFold3 and AlphaPulldown2 models of the ORP11-SAC1 complex embedded in ER-like flat bilayers generated using CHARMM-GUI. Residues within 3.5 Å of the upper membrane leaflet are coloured green, indicating potential membrane contacts. Circled is the helix in SAC1 that we report to be amphipathic. **B**. The same models coloured by domain architecture to highlight structural differences. In the AlphaFold3 model, the PH domain (in yellow) of ORP11 is positioned closer to the membrane, whereas in the AlphaPulldown2 model, the PH domain interacts directly with the SAC domain of SAC1.

We examined the α-helices in SAC1 that lie near the upper leaflet of the bilayer to assess their amphipathic character using HeliQuest ^58^. Only the segment “FLAMLNHVLNVDGFYFST” (residues 119-135) shows notable hydrophobicity (0.77). This region (marked with a circle in **Fig. 5A**), is located very close to the membrane and may also contribute to the stabilisation of the ORP-SAC1 pair complex (**Fig. S5**). However, AlphaFold predicts that about half of these residues form a curved loop extending away from the membrane, suggesting that this helix may simply require model refinement in AlphaFold rather than forming a stable amphipathic region above the curved bilayer.

We later considered whether pathogenic mutations in ORP11 might target the SAC1 interaction interface, potentially explaining a molecular mechanism underlying disease. In particular, OMIM (https://omim.org/) reports a case describing two brothers from consanguineous Turkish parents with a severe neurodegenerative phenotype, including psychomotor retardation, ataxia, and demyelinating leukodystrophy. Genetic analysis revealed a homozygous missense mutation in the *POLR1A* gene, and an R171W substitution in *OSBPL11*, the gene encoding ORP11 ^59^. Although initially intriguing, the R171W mutation lies in a disordered region of ORP11, between the PH domain and the coiled-coil domain and is distant from the SAC1 interface in our *in silico* models of the ORP11-SAC1 complex. Therefore, a direct disruption of the interaction is unlikely. However, we cannot exclude the possibility that this mutation affects overall protein folding, stability, or dimerisation, which may indirectly impair ORP11 function. No disease-associated mutations have been reported for SAC1 or ORP11’s closest paralogues, ORP9 and ORP10.

### 3.4 Higher-order ORP dimeric assemblies explored

Following our analysis of ORP-phosphatase interactions, we turned our attention to the possibility of higher-order assemblies involving ORP9, ORP10 and ORP11. This direction was motivated by a recent study reporting that ORP9 and ORP11 form heterodimers ^13^. We sought to determine whether state-of-the-art structure prediction tools could recapitulate this dimer interface, whether such a dimer could be predicted with any degree of confidence, and whether SAC1 might be incorporated into these higher-order assemblies.

We summarise here the results from AlphaFold3 and AlphaFold2-Multimer predictions for the ORP10-ORP11 dimer (**Figs. S6-S7**). Overall, confidence scores were low across both tools. Importantly, the predicted dimer interface differed substantially between AlphaFold3 and AlphaFold2, highlighting a lack of convergence. The AlphaFold3 model positioned the PH domains of both ORPs in close proximity and suggested a potential coiled-coil-mediated interaction. The PAE heatmap showed medium-confidence contacts at this interface, but corresponding pLDDT scores were uniformly low (**Fig. 4C**), suggesting significant uncertainty or potential disorder in this region. The AlphaFold2 model also showed little overall confidence, with the actifpTM score clearly reflecting low interface reliability (**Fig. S7**).

Next, we tested whether SAC1 could be integrated into this putative ORP10-ORP11 dimer. Adding a single SAC1 molecule in the AlphaFold3 prediction resulted in severe steric clashes between the ORD domains of ORP10 and ORP11 (**Fig. S8A**), ruling out this configuration. In contrast, AlphaFold2-Multimer was able to place SAC1 without clashes (**Figs. S8B-C**). This model showed one ORD domain from ORP10 engaging SAC1 through the interface we have consistently seen in previous models, while the second ORD domain from ORP11 approached SAC1 from a lateral angle, forming additional contacts. The actifpTM score here indicated moderate to high confidence in this asymmetric interaction. Still, this model should be interpreted with caution, as only AlphaFold2 predicted it without generating steric conflicts, and the interface was not replicated by AlphaFold3.

Finally, we examined a tetrameric configuration where two SAC1 molecules bind an ORP10-ORP11 dimer, testing whether SAC1 might interact with each ORP separately. The resulting models, shown in **Fig. S9**, failed to support this architecture. Confidence scores dropped again, and both AlphaFold3 and AlphaFold2 produced structurally incompatible configurations. In AlphaFold3 and AlphaFold2, the TM domains of SAC1 were misoriented relative to a hypothetical membrane bilayer. PAE heatmaps highlighted widespread uncertainty in domain positioning and overall spatial arrangement, undermining the plausibility of this tetrameric complex.

We next turned to the ORP9-ORP11 pair, which has been reported to form dimers ^13^. AlphaFold3 consistently predicted higher ipSAE scores than weighted ipTM+pTM scores, regardless of SAC1 inclusion (**Fig. S10**). The model suggests that dimerisation occurs through their coiled-coil domains. However, when a single SAC1 molecule is added, the luminal side of its TM domain interacts with residues on ORP9’s ORD domain: a biologically implausible scenario, as this region should be membrane-embedded (**Fig. S10B**). Interestingly, when two SAC1 chains are included to form a tetramer with ORP9-ORP11 (**Fig. S10C**), the complex becomes symmetrical and SAC1’s TM domains dimerise. Despite this, the pLDDT and weighted ipTM+pTM scores drop compared to the other structures, though ipSAE and pDockQ remain above 0.5.

In all AlphaFold3 models, the PAE plots fail to show confident alignment between ORP9 and ORP11. This suggests that the structural models do not strongly support a stable, direct interaction between the two proteins, despite their known tendency to dimerise. In **Fig. S11**, we modelled the same complexes using AlphaFold2-Multimer. The actifpTM scores are all above 0.9, indicating high confidence. Still, SAC1’s TM domains are misoriented in one of our multi chain structures (**Fig. S11C**): they are not perpendicular to the plane of the membrane. Once again, the PAE heatmap points to uncertainty in the relative positioning of ORP9 and ORP11, reinforcing the point that the PAE never shows any kind of confidence and that the interaction between them could not be modelled confidently. Given the flexibility between domain regions in ORP9 and ORP11, conformational shifts could in principle allow SAC1 to align correctly, but this isn’t captured in the current model.

## 4 Discussion

Interpreting predicted protein-protein interactions requires more than structural plausibility, it demands a biologically grounded framework that filters out artifacts and elevates meaningful contacts. In this study, we only consider an interaction to be confidently predicted when multiple independent lines of evidence align. High interface quality, measured primarily through actifpTM scores and supported cautiously by ipSAE, is a first requirement. Good scores alone are insufficient. Predictions must show compatible subcellular localisation. Interactions between ER- and cytosol-localised proteins are plausible for peripheral or integral ER proteins only if the interaction surface maps to the cytoplasmic region; luminal ER proteins, by contrast, are topologically isolated and unlikely to engage cytosolic partners. This topological consideration is an important factor that is often overlooked but critical for filtering biologically meaningful interactions. Structural plausibility is equally critical, particularly for membrane-associated complexes: the predicted interface must not violate known membrane topologies or orientations. Finally, consistency across models between AlphaFold2 and AlphaFold3, and/or among homologues adds crucial support. pLDDT and congruent interface geometries across models indicate structural reliability. By this stringent standard, only a minority of predictions are accepted. The ORP11-SAC1 complex is one such case, satisfying all criteria and exhibiting high-confidence interface geometry across the methods used.

These criteria are not arbitrarily conservative; they reflect fundamental architectural and methodological differences between AlphaFold versions and pipelines. AlphaPulldown2 builds on AlphaFold2-Multimer, whose Evoformer module extracts co-evolutionary information from multiple sequence alignments (MSAs) and refines pairwise residue representations through transformer-based iterations. The Evoformer is effective when deep MSAs are available but suffers in sparse evolutionary landscapes ^60^. AlphaFold3 is very different. Its multimodal diffusion-based architecture accepts not only sequence, MSA input and structural templates, but ligands, and ions ^34^. As a result, AlphaFold3 can succeed where AlphaFold2 and AlphaPulldown2 fail, but also diverge at times, producing markedly different interfaces, loop conformations, and subunit orientations even from the same input sequences.

### 4.1 Assessing interactions with new generation confidence scores

Given the considerations above, it is important to note that model confidence scores from different methods (i.e. ipSAE vs actifpTM) can vary substantially and do not always agree. While the scores all operate on a nominal 0-1 scale, they are not always directly comparable. Independent studies have established empirical thresholds for model confidence for one of the metrics we use. pDockQ scores above 0.23 are predictive of likely interactions ^36,42^, among which scores of ≥0.80 indicate high-accuracy models, 0.49-0.79 correspond to medium accuracy, and 0.23-0.48 are considered acceptable ^37^. Our best models in this study fall within the 0.3-0.5 range, consistent with acceptable to medium-quality predictions. While pDockQ is not a definitive standard for model assessment, and arguably no such gold-standard metric exists yet, it still provides a useful frame of reference.

The ipSAE score is another key metric we used, and it is not limited to interface residues ^27^. Instead, it is based on all inter-chain residue pairs with PAE(i,j) values below the chosen PAE cutoff. The interface distance parameter in the implementation is used only to report the number of residues meeting both the PAE and distance criteria, but it does not contribute to the ipSAE value itself. The score defaults to zero only when there are no inter-chain residue pairs below the cutoff, i.e., when AlphaFold predicts no confident relative positioning of residues across chains. This feature differentiates ipSAE from ipTM and related scores, which rarely approach zero because they consider all residue pairs regardless of confidence.

The ipSAE score has only recently been introduced and, while increasingly used ^61-63^, has not yet been widely benchmarked by groups independent of its developers. In its original report, benchmarking on sets of true and false complexes showed that almost all false complexes scored below 0.2, while true complexes with correct structural models generally scored above 0.5, with few intermediate values ^27^. In our dataset, the highest ipSAE values (≥0.6, for the ipSA_d0dom score employed here) were observed in complexes involving the phosphatidylinositol 3- and 4-phosphatase SAC1, placing them above the benchmarked threshold.

### 4.2 Biological implications of ORP-PIP interactions

A central point of this study is that structure prediction alone, especially from a single method, is often insufficient to infer functional interactions. To strengthen our findings, we incorporated subcellular localisation data from the Human Cell Map ^52^, Prolocate ^53^ and Map of the Cell ^54^ databases. The Human Cell Map, based on BioID rather than fractionation, captures proximity relationships instead of steady-state distributions. This can place proteins at transient or peripheral contact sites and, in our case, highlights ORP2 and ORP8 at the nuclear outer membrane-ER network, representing a potentially new localisation (**Fig. S12A**). ORP3 and INPPL1 (SHIP2) are assigned to the PM, while a substantial fraction of the remaining proteins of interest is placed at early recycling endosomes (**Fig. S12A**). This likely reflects the proximity-labelling approach of the Human Cell Map, where vesicular trafficking hubs are frequently captured because of the high connectivity of endosomal compartments ^52^.

In contrast, Map of the Cell annotations are less comprehensive, with many proteins missing localisation assignments. Only SAC1, ORP3, and ORP10 are assigned to the ER, and ORP9 together with INPP5A are placed at the PM (**Fig. S12B**). In this scenario, INPP5A could act as a receiver or processor of lipids transferred across the junction, helping maintain membrane lipid balance, potentially a role SAC1 fulfils considering its models confidently with ORP9 and ORP11 across multiple tools and that these protein pairs structurally align strongly. Overall, the discrepancies across datasets arise from the methodological contrast between proximity-labelling (Human Cell Map) and biochemical fractionation (Map of the Cell). As a result, each method can highlight different protein localisations.

Here, we review examples of Prolocate fractionation data from rat liver that illustrate how subcellular localisation can inform potential protein-protein interactions: (1) SAC1 and INPP5K are reported to localise at least partially to the ER, supporting the feasibility of their interaction with ER-tethered ORPs; (2) ORP11 and SAC1 are found in overlapping ER- and cytosol-enriched fractions; (3) INPP5K and SYNJ2 show partial localisation to ER-PM junctions, possibly through interactions with less well-characterised ER-anchored partners beyond the canonical recruiter ARL6IP1 ^10^; (4) most ORPs are cytosolic, consistent with their role as lipid shuttles, but exceptions include ORP2 at the Golgi and ORP5 and ORP8 at the ER (**Fig. S12C**).

When considering cellular compartments, it is also useful to examine the detailed dynamics at MCSs. Although membrane association might seem incompatible with a classical shuttle mechanism, it remains theoretically possible for membrane-anchored ORPs to mediate lipid exchange across short intermembrane distances, such as those at MCS (∼10 nm) ^3,4^. If their lipid-binding domains contain sufficiently long flexible loops or linkers, they might even bridge wider gaps. In some mammalian cells, ER–plasma membrane MCSs have been observed to reach up to 25 nm ^64^. In this view, even ER- or Golgi-localised ORPs might still operate in a shuttling mode, delivering lipids to an adjacent membrane without requiring full dissociation into the cytosol.

The biological relevance of the ORP11-SAC1 interaction is further underscored by the absence of FFAT motifs in ORP11, ORP10, ORP5, and ORP8. FFAT motifs typically mediate ER anchoring via interaction with VAPA/B ^14,18,19^. In these FFAT-lacking ORPs, SAC1 or other ER-resident partners may substitute as recruitment factors. This mechanism is consistent with models proposed for OSBP- and ORP5/8-mediated transport, where PI4P is transferred to the ER and dephosphorylated by SAC1 to drive lipid exchange against a concentration gradient ^12,21^. In these models discussed by other groups, SAC1 acts as a functionally coupled but physically separate enzyme; in contrast, our findings suggest a direct physical interaction between SAC1 and ORP11 that we expect is necessary in order to deliver lipid or receive from the ORD domain ^15^.

Accessory proteins such as SAC1 may play dual roles: facilitating ORP recruitment to ER-associated membrane contact sites and potentially supporting lipid exchange. SAC1 could act downstream of lipid transfer by converting incoming PI4P to PI, thereby sustaining lipid gradients and directional flow. Strict coupling between lipid transfer and SAC1’s enzymatic activity may not be required, mere proximity to the site of PI4P delivery could suffice for efficient exchange. A key question to be investigated is whether ORPs preferentially bind phosphorylated or dephosphorylated lipids, which would clarify their functional interplay. We suggest that ORP11 interacts with SAC1 only as a monomer and not as an ORP9-ORP11 heterodimer. In this model, SAC1 resides at the ER while the ORP11 ORD domain transiently shuttles between membranes to pick up and deliver lipids. An unresolved issue is whether lipid binding alone is enough to trigger ORD translocation to the opposing membrane, or whether a coordinated mechanism exists in which the PH domain remains bound at the PM or Golgi while the ORD domain engages the ER, alternating positions to shuttle lipids across (**Fig. 6**).

**Fig. 6.**
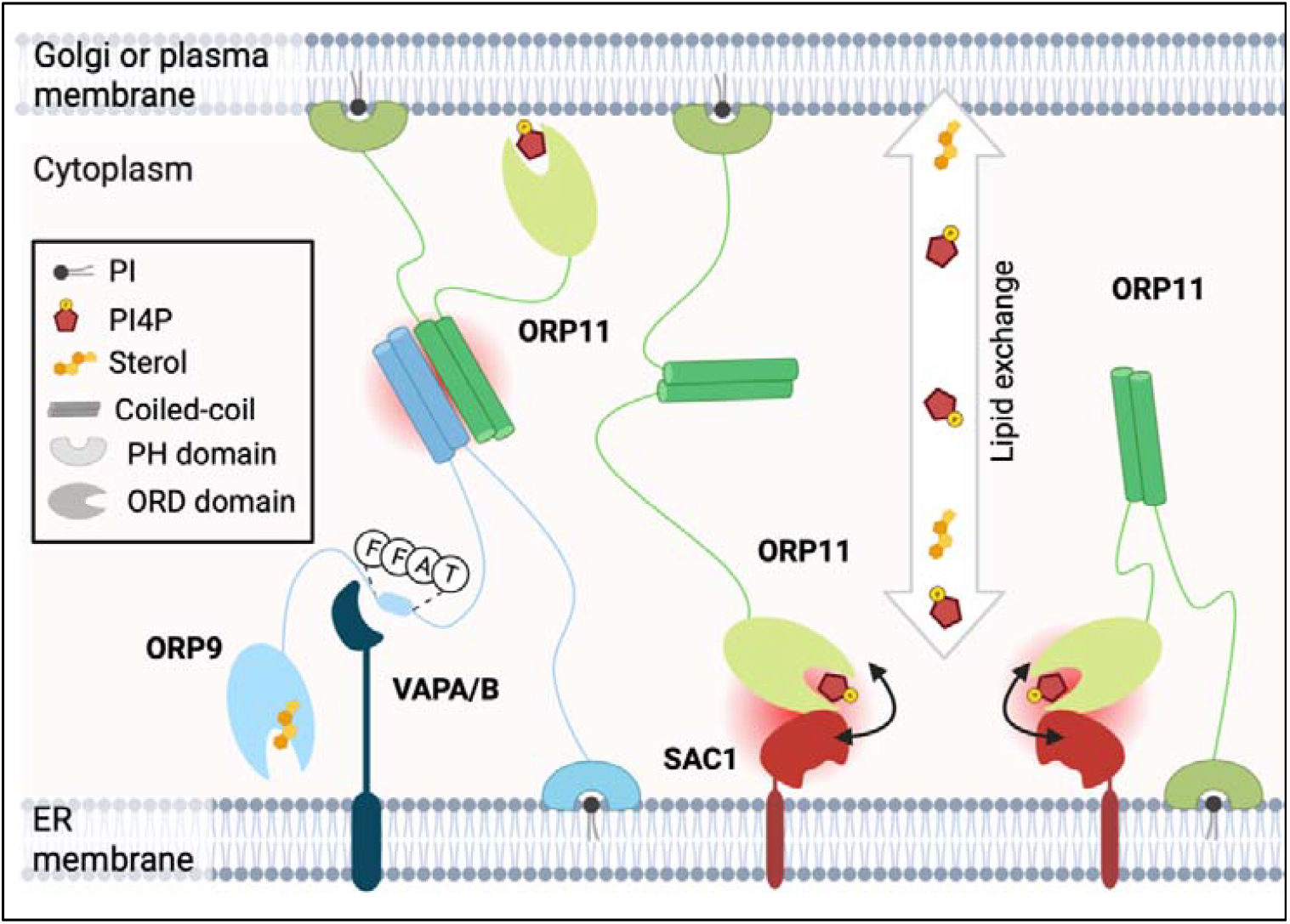
SAC1 recruits a single molecule of ORP11 to the ER. Literature shows that ORPs localise to ER-PM and ER-Golgi contact sites. The ORP9-ORP11 heterodimer, formed through their coiled-coil domains, bridges these junctions. If bound to VAPA/B via its FFAT motif, ORP9 is anchored to the ER, while ORP11 can occupy the PM membrane surface (shown on the left). If ORP11 is not forming a dimer with ORP9 and acts as a single molecule, we propose that it is recruited by SAC1 to the ER. The ORP11 PH domain could still span the contact site distance and anchor at the PM (shown in the centre), or more simply sit at the ER cytosolic leaflet (shown on the right). SAC1 could: (1) modulate ORP11’s lipid transfer activity by dephosphorylating cargo either before export from the ER when it acts as the donor membrane, or after import when it serves as the acceptor membrane; (2) contribute to lipid loading or unloading onto the ORP11 ORD domain spanning the junction. Red circles indicate protein-protein interactions.

Our results align with the concept that SAC1 activity is tightly regulated by ORP family proteins at MCSs, as previously suggested in yeast and discussed here. In *Saccharomyces cerevisiae*, SAC1 controls PI4P levels across the ER, Golgi, and PM, and its activity requires the oxysterol-binding homology (Osh) proteins. Notably, Osh3 localises to ER-PM contact sites in a PI4P-dependent manner, and both *in vitro* and microsomal assays demonstrate that Osh proteins directly stimulate yeast SAC1 phosphatase activity ^25^. Although attempts to detect a stable complex between the purified yeast SAC1 domain and Osh3 or Osh4 ORDs *in vitro* were unsuccessful, co-immunoprecipitation experiments indicated that only a small fraction of SAC1 can interact with Osh proteins in cells ^25^, consistent with SAC1’s approximately 20-fold higher abundance compared to Osh7 ^65^. These findings suggest a conserved principle whereby ORPs can act both as sensors of membrane lipid composition and as activators of SAC1, enabling localised PI4P turnover and facilitating directional lipid exchange ^23,25,66^.

The subcellular localisation data lend weight to our structural predictions, though they have not been directly validated. Proteomic data, particularly from fractionation and imaging studies, offer complementary but often imperfect perspectives. Conventional fluorescence microscopy offers limited resolution, around 200 nm, which corresponds roughly to the size of a large vesicle, far larger than the nanometre-scale gaps at MCSs ^67,68^. Super-resolution imaging methods (i.e. STEDM, PALM) can achieve finer spatial resolution and may provide more informative localisation data at these small scales ^67^. Proximity labelling techniques like BioID leverage covalent tagging of neighbouring proteins in living human cells and can generate high-resolution localisation maps in human cells, as exemplified by the Human Cell Map project ^52^. In contrast, subcellular fractionation separates cellular compartments biochemically and infers protein localisation based on distribution across fractions, providing a global overview.

To analyse subcellular localisation, we used two major databases reporting data from fractionation assays: Prolocate ^53^ and Map of the Cell ^54^. The Prolocate dataset derives from two experiments on rat liver, one from fasted and one from fed animals (**Fig. S12C**). The data from Prolocate comes from two experiments: one conducted on fasted rats and one on fed rats. As a result, experiment A, which was performed on fasted rats, does not reflect conditions that are fully comparable to those obtained under healthy and normal conditions. More work is needed to confirm that both ORP11 and SAC1 co-reside and function at the same contact sites in human cells, ideally with high resolution imaging and biochemical interaction experiments. Nonetheless, this approach offers a more reliable localisation framework than antibody-dependent databases like the Human Protein Atlas, where issues of non-specificity often limit interpretability.

The Map of the Cell database was created using HeLa cells rather than rat liver. Like Prolocate, it is based on subcellular fractionation but in a human cancer cell line. HeLa cells have an abnormal number of chromosomes (aneuploidy) and extensive genomic rearrangements. This genomic instability alters gene regulation and can directly impact the abundance and localisation of many proteins and the stoichiometry of protein complexes compared to non-cancerous cells. As a result, protein localisation in HeLa cells may differ from that observed in rat liver cells due to a combination of chromosomal abnormalities and regulatory changes. For example, while the Prolocate dataset reports the largest fractions of ORP9, ORP10, and ORP11 at lysosomes/peroxisomes, cytosol, and cytosol, respectively, the Map of the Cell dataset instead assigns them to the PM (ORP9), ER (ORP10), and Golgi apparatus (ORP11).

Our overarching goal was to situate AlphaFold-based structural predictions within a robust biological context. After highlighting high-confidence complexes such as ORP11-SAC1, we examined the rest of the dataset. Beyond the ORP9, ORP10, and ORP11 trio, which provide strong interface scores, other clusters of protein pairs, such as ORP5-SAC1, ORP8-SAC1, ORP6-SAC1 and ORP7-SAC1, structurally align well. These interactions may not be strong enough to form stable complexes, but they merit further investigation. In fact, they may operate through coincidence detection, a principle well-recognised in cell biology. Coincidence detection describes systems in which biological outcomes only arise when multiple conditions are simultaneously satisfied ^69,70^. At MCSs, ORPs and phosphatases co-residing transiently may create the potential for interactions that are weak individually but functionally relevant in the right context. Conversely, the lowest-confidence predictions, which lack structural alignment with other protein pairs, suggest that direct physical interactions are unlikely or too transient to capture computationally. In these cases, interfaces may be too small or weak to model reliably.

### 4.3 AlphaFold3 lipid binding predictions and limitations

To further evaluate the biochemical plausibility of the ORP11-SAC1 interaction, we modelled the complex in AlphaFold3, including molecules of palmitic acid (PLM). When modelling ORP11-SAC1 with lipid in AlphaFold3, one of the PLM molecules is very closely positioned near the catalytic site of SAC1, highlighted in **Fig. S4A**, which was shown to be crucial since a mutation in position 389 completely abrogates its activity *in vitro* ^12^. We employed AlphaFold3 to model the complex with larger numbers of lipids, representative of bulk phospholipids like in a membrane bilayer. Our approach was motivated by the observation that including lipid ligands to mimic a membrane environment can improve confidence and accuracy in predicting membrane-associated protein complexes ^71^. With recent upgrades in AlphaFold3’s multimodal architecture, limited modelling of protein-small molecule interactions has become possible, which we aimed to exploit.

It is important to note that PLM is not the physiological ligand for either SAC1 or ORP11. SAC1 specifically dephosphorylates PI4P, while ORP11 is involved in the exchange of PI4P with other lipids such as phosphatidylserine or cholesterol. As such, PLM serves only as a proxy for hydrophobic tail occupancy and cannot reproduce the electrostatic or headgroup-specific interactions that govern true lipid recognition.

### 4.4 Comparative analysis of predicted protein-protein interfaces

We applied an interface lDDT score cutoff of 0.8 for structural alignment in FoldSeek-Multimer. This stringent threshold was chosen because, after excluding ORP protein regions not at the interface, and therefore not part of the ORD domain, we expected high structural similarity between models. Trimming one chain from each protein dimer effectively removes many residues or coincidental contacts in regions like the PH and coiled-coil domains that do not contribute to a consensus interface.

Using this 0.8 interface lDDT cutoff, we identified a coherent subcluster of SAC1-bound ORPs. We used this high threshold because after removing ORP protein regions that are not at the interface, we can expect the structural similarity of the interacting residues to be very high. This way by trimming down one of the chains of each protein dimer, we are filtering out already a lot of residues or coincidental contact points in the PH domain and coiled-coil domain that are not relevant to a consensus interface. This forms a cluster including AlphaPulldown2 and AlphaFold3 models, all displaying moderate interface metrics (ipTM+pTM > 0.5, pDockQ > 0.23, actifpTM > 0.7) and closely aligned geometries at the SAC1-ORD interface. Notably, this comprises ORP10-SAC1, and ORP9-SAC1, which show striking similarity to ORP11-SAC1, with surprisingly strong alignment consistency between AlphaPulldown2 and AlphaFold3 predictions (**Fig. S2**).

The FoldSeek-Multimer methodology is sensitive to broader conformational differences, particularly when large mobile domains like the PH or coiled-coil regions adopt variable orientations or display disorder. In the case of ORP10-SAC1, AlphaFold3 and AlphaPulldown both predict similar interface contacts, yet the ORD domain adopts a different angle relative to SAC1 between models. This change likely contributes to a lower interface similarity score. While this could reflect a real difference in predicted complex geometry, several other factors might play a role ^35^. First, FoldSeek-Multimer aligns structures globally before assessing local interfaces, meaning even small shifts in domain orientation can misalign conserved binding regions. Second, FoldSeek-Multimer evaluates entire structures chain-to-chain and does not align them by selecting interface residues and superimposing them, so well-conserved binding regions can be underweighted. Third, modest differences in sidechain positioning or flexible loops at the interface which do not affect apparent interface topology can still lower lDDT scores.

In contrast, ORP6-SAC1, ORP7-SAC1, ORP5-SAC1, and ORP8-SAC1 show consistent binding interfaces across AlphaFold2 and AlphaFold3 models, results that are not seen in the full-length models being clustered but only when the ORPs were trimmed down to the ORD domain. The predicted contact between SAC1’s TM domain and ORP coiled-coil domains (i.e. in ORP4, 6, 7) is biologically unlikely, reflecting false dimerisation due to similar helix packing.

To investigate the artefactual TM-coiled-coil interaction a little further, we modelled ORP4-SAC1, ORP6-SAC1, ORP7-SAC1 with 25 palmitic acid molecules. Lipid coverage disrupted binding at the TM surface, but the TM helices’ luminal side engaged residues in ORP, which is not biologically plausible since ORPs cannot cross membranes and should be modelled only above the TM plane. Finally, removing SAC1’s TM domain and modelling it with full-length ORPs caused drastic orientation changes, with PH domains blocking the SAC interface, contradicting the known lipid-binding role of these domains.

### 4.5 ORP dimerisation

Finally, we explored the assembly of higher-order ORP complexes, specifically focusing on experimentally-proven ORP9-ORP11 and potential ORP10-ORP11 heterodimers. Our attempts to model a similar dimer between ORP10 and ORP11 did not yield confident structural predictions. This interest stems from recent findings showing that ORP9 and ORP11 dimerise to localise efficiently at ER-Golgi contact sites ^13^. Dimerisation occurs through coiled-coil regions located between the PH and ORD domains ^17^. Previous analyses using AlphaFold-Multimer and PCOILS indicate that both ORP9 and ORP11 possess two α-helices in this region that form a stable dimer through coiled-coil interactions ^13^. Using a combination of AlphaFold2-Multimer and AlphaFold3, we successfully modelled two plausible conformations of the experimentally validated ORP9-ORP11 dimer, including scenarios where SAC1 is simultaneously engaged at the interface.

Our findings loosely align with experimental data: co-immunoprecipitation experiments demonstrate that ORP11 interacts with VAPA only in the presence of ORP9, and this interaction is abolished in cells lacking ORP9 or expressing a VAPA mutant deficient in FFAT binding ^13^. This confirms that ORP11, although lacking a FFAT motif of its own, can access ER tethers indirectly via dimerisation with FFAT-containing ORPs. The PH domains of both ORP9 and ORP11 mediate Golgi localisation through interaction with phosphatidylinositol phosphates ^17^, anchoring the complex at ER-Golgi contact sites. Finally, we did not model ORP paralogues forming homodimers, as experimental evidence from co-immunoprecipitation assays shows that ORP9 and ORP10 do not form homodimers ^72,73^.

Taken together, our results reveal a structurally and biologically coherent model in which SAC1 not only interacts directly with ORP11 but may also serve as a general hub for recruiting FFAT-deficient ORPs to ER membranes. Given that ORP11 forms heterodimers with ORP9, this raises the intriguing possibility that SAC1, ORP11, and ORP9 form a trimeric complex at MCSs. These interactions exemplify the modularity of the ORP family and underscore the value of high-confidence in silico predictions guided by proteomic and cellular context. They suggest that membrane contact sites are shaped not just by strong, stoichiometric interactions but by a network of opportunistic, context-dependent associations, many of which may be mediated by coincidence detection mechanisms. In this light, structure prediction becomes a way to map dynamic interfaces central to organelle communication other than just to identify stable complexes.

## 5 Conclusion

Our study demonstrates how machine learning-based structural prediction can be integrated with proteomic, subcellular localisation, and evolutionary data to uncover biologically relevant interactions at membrane contact sites. By applying AlphaPulldown2 and AlphaFold3 at scale, and introducing confidence metrics such as actifpTM, interface lDDT, and FoldSeek-Multimer clustering, we mapped a structural interaction landscape between lipid transfer proteins and phosphoinositide phosphatases. Among these, the ORP11-SAC1 complex stands out as a high-confidence interaction that satisfies structural, biochemical, and contextual criteria. Our findings also suggest that SAC1 may act not only as a phosphatase but as a recruiter or stabiliser for FFAT-lacking ORPs at ER-associated sites. Attempts to refine models with lipid molecules highlight the potential and current limits of applying structure prediction tools to membrane-bound and -aware protein-lipid systems. Altogether, this work provides a blueprint for how AI-guided structure prediction, when paired with biological reasoning, can advance our understanding of dynamic, low-affinity, and context-dependent interactions that are crucial at organelle contact sites.

## Author Contributions

**Dall’Armellina, F**: Software, Validation, Formal analysis, Investigation, Data Curation, Writing - Original Draft, Visualisation. **Urbé, S**: Supervision, Resources, Review and Editing. **Rigden, DJ:** Supervision, Resources, Review and Editing, Methodology, Project administration.

## Acknowledgments

Funding was provided by the Institute of Systems, Molecular and Integrative Biology (ISMIB) at the University of Liverpool, supported by the Douglas Endowment Fund. We are grateful to Prof. Patrick A. Eyers, Prof. Sonia Rocha, and Prof. Michael J. Clague for their support of this project. We also thank Prof. Roland L. Dunbrack Jr. for providing feedback on the ipSAE and related scores. We thanks Dr. Jools Wills (Research Complex at Harwell) for computational support. Additional thanks go to Dr. Shahram Mesdaghi for discussions and feedback on AlphaPulldown, and Dr. Emily Johnson for input on FoldSeek-Multimer and network clustering; both are members of the Computational Biology Facility (CBF) within the Liverpool Shared Research Facilities.

## Data Availability Statement

Models were reposited in Figshare, while the SAC1-ORP dimer cluster is available on ModelArchive ^74^.

## Supplementary Information

**Figs. S1-S12**

**Fig. S1.**
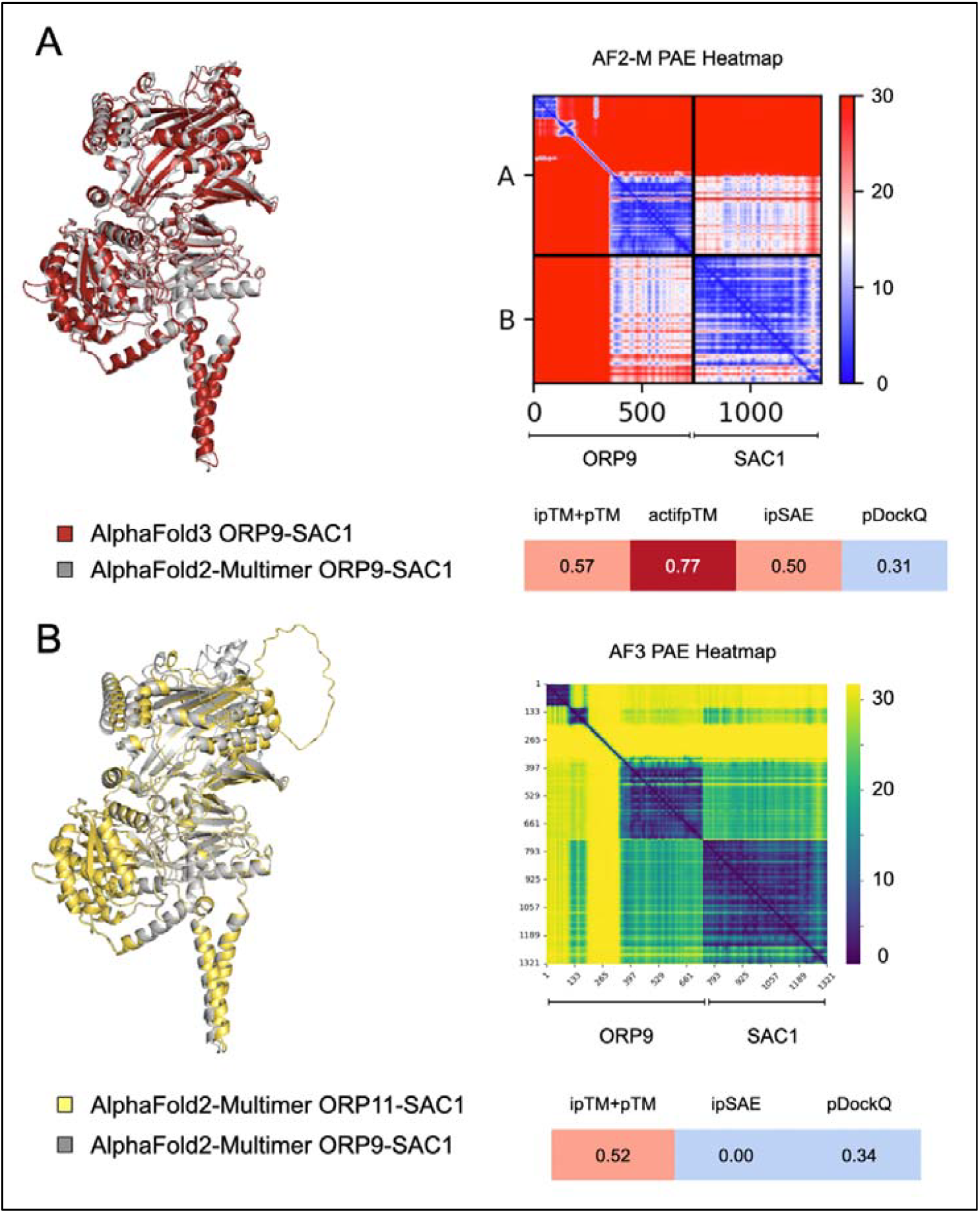
**A.** AlphaFold3 and AlphaFold2-Multimer ORP9-SAC1 complexes superimposed. **B**. AlphaFold2-Multimer ORP9-SAC1 and ORP11-SAC1 complexes superimposed (centre). PAE heatmaps and interaction confidence scores for the AlphaFold3 and AlphaFold2-Multimer ORP9-SAC1 models on the right. “ipTM+pTM” represents the weighted confidence score, calculated as 0.8 × ipTM + 0.2 × pTM. “ipSAE” represents the ipSAE_d0dom score.

**Fig. S2.**
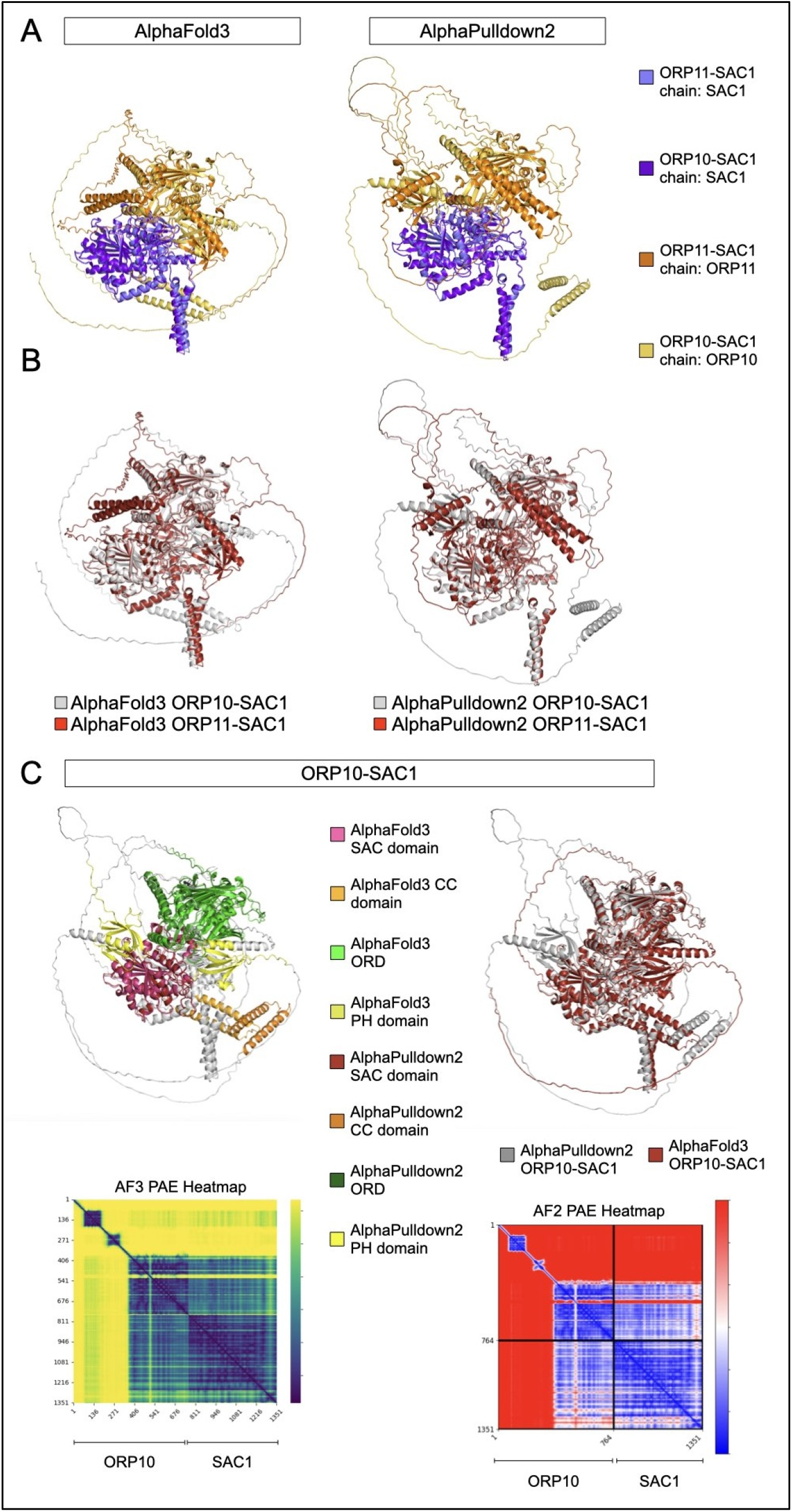
**A.** Structural models of the ORP10-SAC1 and ORP11-SAC1 complexes generated using AlphaFold3 and AlphaPulldown2, superimposed and visualized in PyMOL. Models are shown with chain-based colouring to distinguish individual protein components. **B**. Same structural models as in (A) but shown with colouring by protein pair object to highlight the interface and interaction regions for each complex. **C**. Focused comparison of ORP10-SAC1 complexes only. AlphaFold3 and AlphaPulldown2 models are shown with domain-based colouring (left) and whole-object colouring (centre). The corresponding PAE heatmaps for the ORP10-SAC1 models from AlphaFold3 and AlphaPulldown2 are shown on the right.

**Fig. S3.**
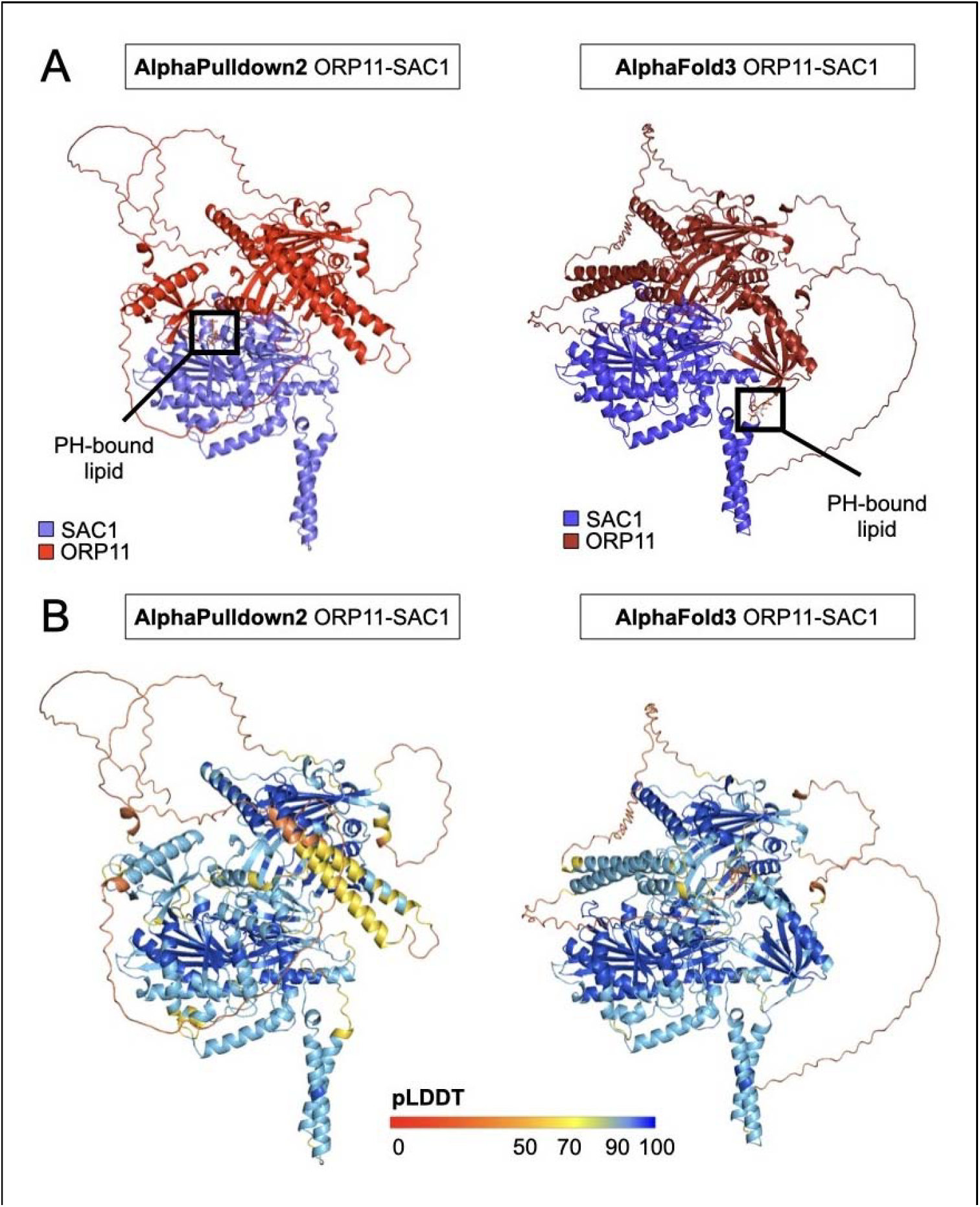
AlphaFold3 and AlphaPulldown2 models of the ORP11-SAC1 complex. **A**. Models are coloured by chain to distinguish SAC1 from ORP11. The PH domain of ArhGAP9 bound to Ins(1,4,5)P_3_ (PDB ID: 2P0D) was superimposed onto each model to assess the orientation of the lipid-binding domain relative to the TM region of SAC1 and the membrane. The AlphaFold3 model positions the PH domain in an orientation more consistent with known lipid binding and membrane topology. **B**. AlphaFold3 and AlphaPulldown2 models of the ORP11-SAC1 complex, coloured by per-residue pLDDT confidence scores. Colour scheme follows the AlphaFold standard (credit: Konstantin Korotkov).

**Fig. S4.**
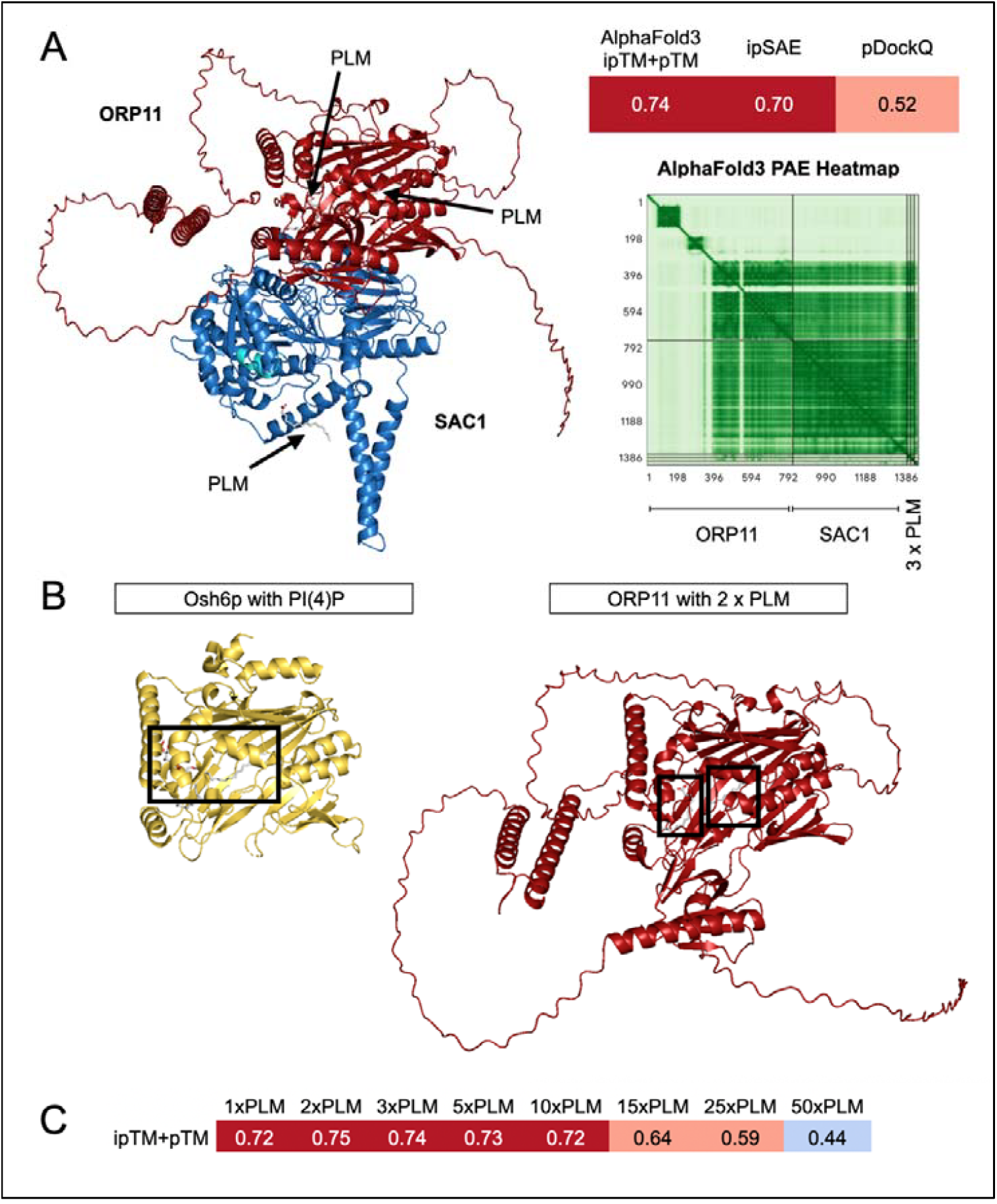
**A**. AlphaFold3 ORP11-SAC1 model in complex with three molecules of palmitic acid (PLM). On the right, results of scoring metrics and PAE heatmap. The catalytic site of SAC1 is highlighted in cyan in the SAC domain. “ipTM+pTM” represents the weighted confidence score, calculated as 0.8 × ipTM + 0.2 × pTM. “ipSAE” represents the ipSAE_d0dom score. **B**. Structure of Osh6p in complex with phosphatidylinositol 4-phosphate or PI(4)P from a published co-crystal (PDB ID: 4PH7). The structure is superimposed to the reference ORP11-SAC1 model in PyMOL. This model is only showing the ORD domain of the AlphaFold3 structure highlighting where two palmitic acid (PLM) molecules were docked. **C**. AlphaFold3 weighted ipTM+pTM scores for modelling ORP11-SAC1 with different numbers of PLM molecules.

**Fig. S5.**
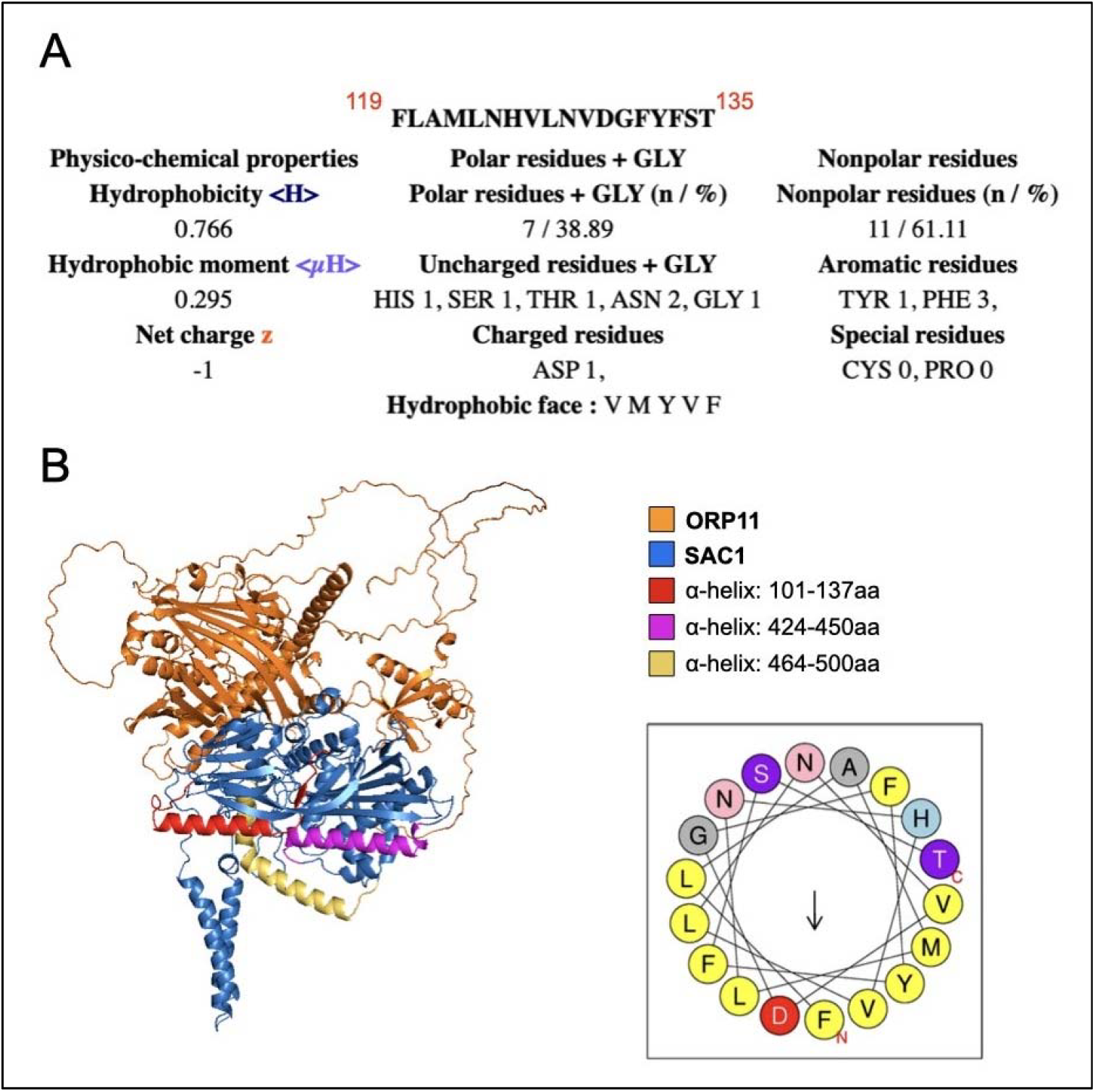
HeliQuest α-helix amphipathicity calculation. **A**. Top-scoring helical segment. **B**. AlphaPulldown2 ORP11-SAC1 model highlighting regions in SAC1 queried to HeliQuest: 101-137, 424-450, 464-500aa. On the right: helical wheel plot of 119-135aa.

**Fig. S6.**
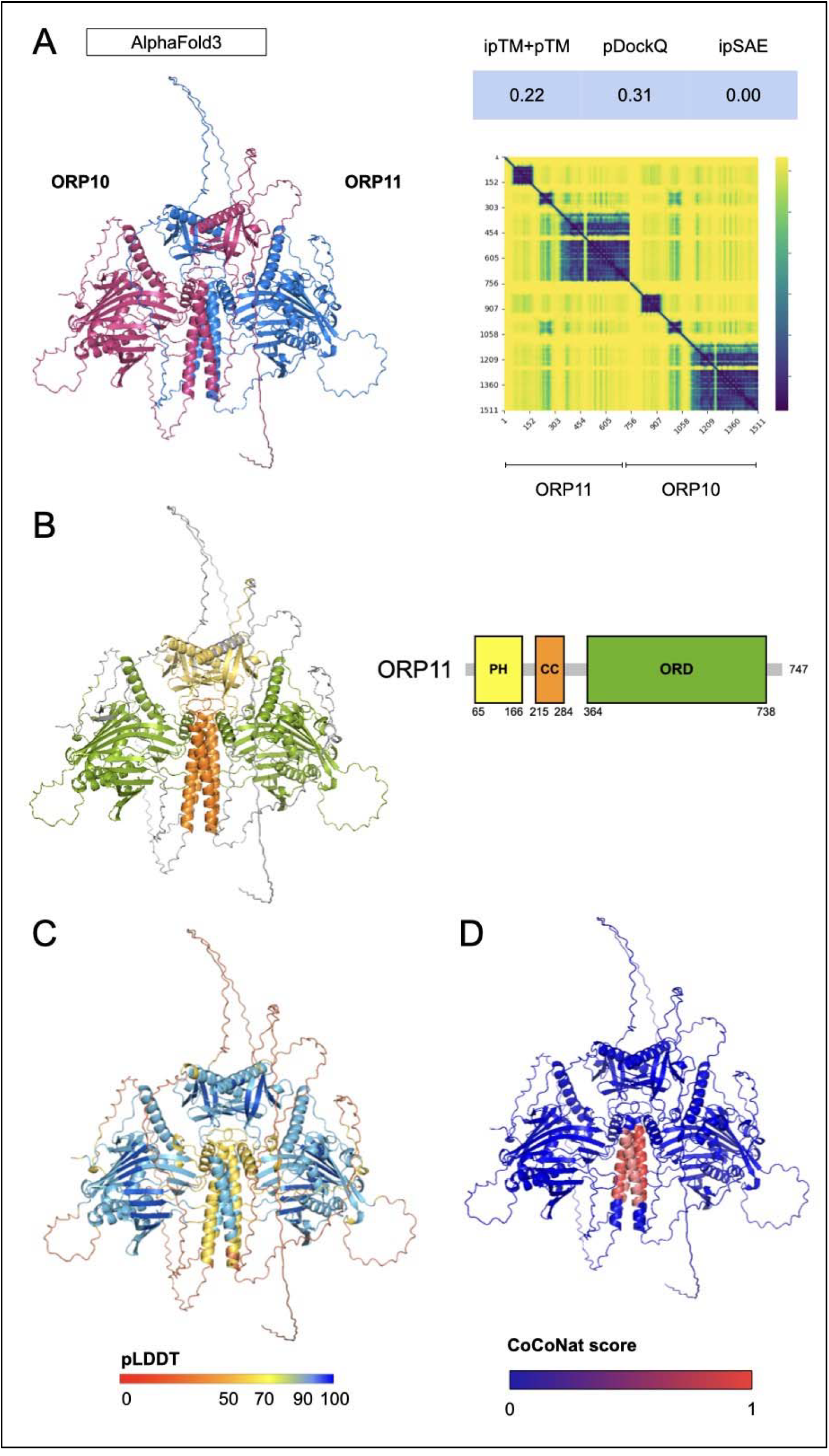
**A**. AlphaFold3 dimer model of ORP10-ORP11 coloured by chain. On the right, scoring metrics and PAE heatmap. “ipTM+pTM” represents the weighted confidence score, calculated as 0.8 × ipTM + 0.2 × pTM. “Interface ipSAE” represents the ipSAE_d0dom score. **B**. Same model coloured by domain architecture. **C**. Coloured by per-residue pLDDT confidence scores. Colour scheme follows the AlphaFold standard (credit: Konstantin Korotkov). **D**. Coloured by per-residue CoCoNat confidence score for predicting coiled-coil domains on a scale 0-1.

**Fig. S7.**
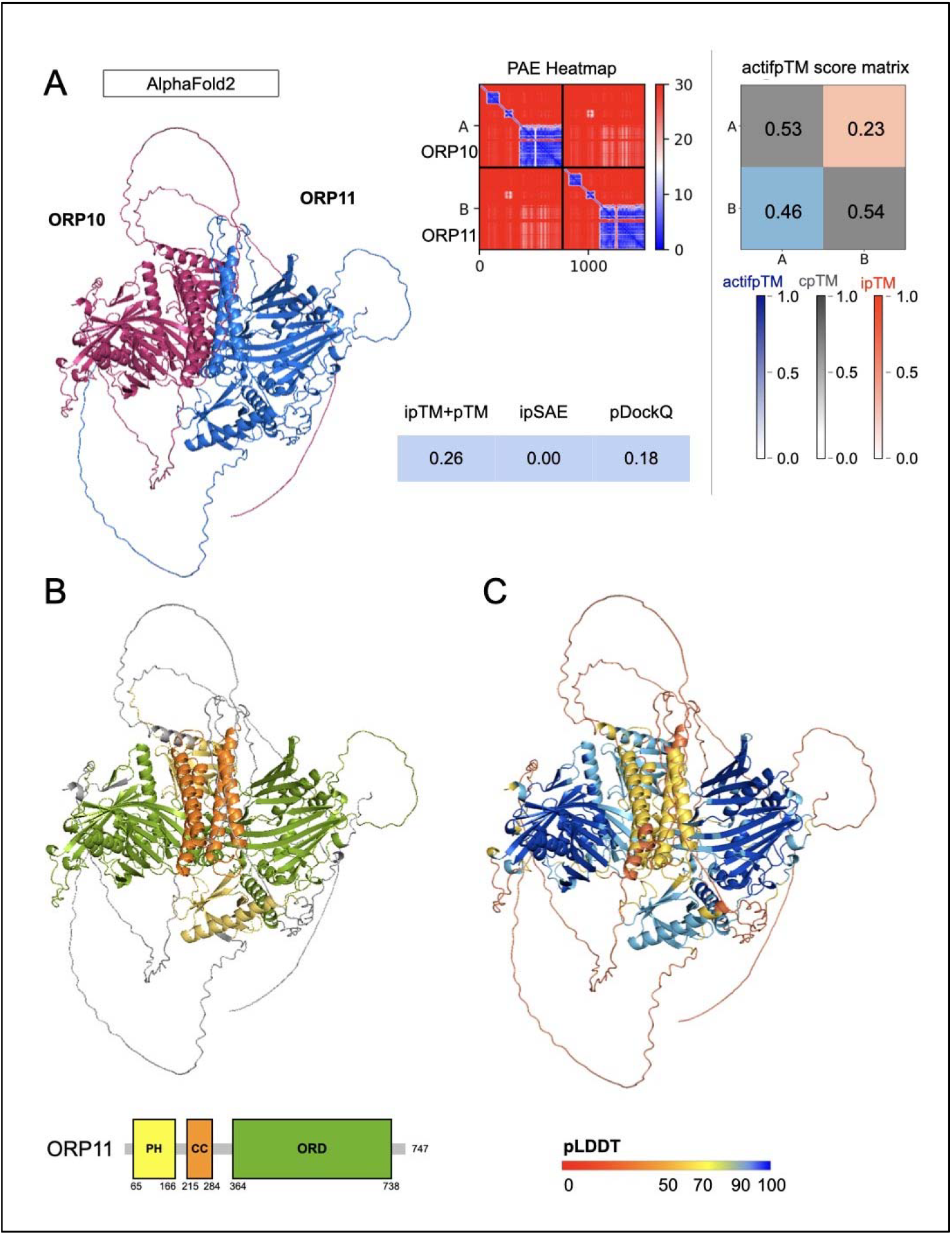
**A**. AlphaFold2 dimer model of ORP10-ORP11 coloured by chain. On the right, scoring metrics and PAE heatmap. “ipTM+pTM” represents the weighted confidence score, calculated as 0.8 × ipTM + 0.2 × pTM. “Interface ipSAE” represents the ipSAE_d0dom score. **B**. Same model coloured by domain architecture. **C**. Coloured by per-residue pLDDT confidence scores. Colour scheme follows the AlphaFold standard (credit: Konstantin Korotkov).

**Fig. S8.**
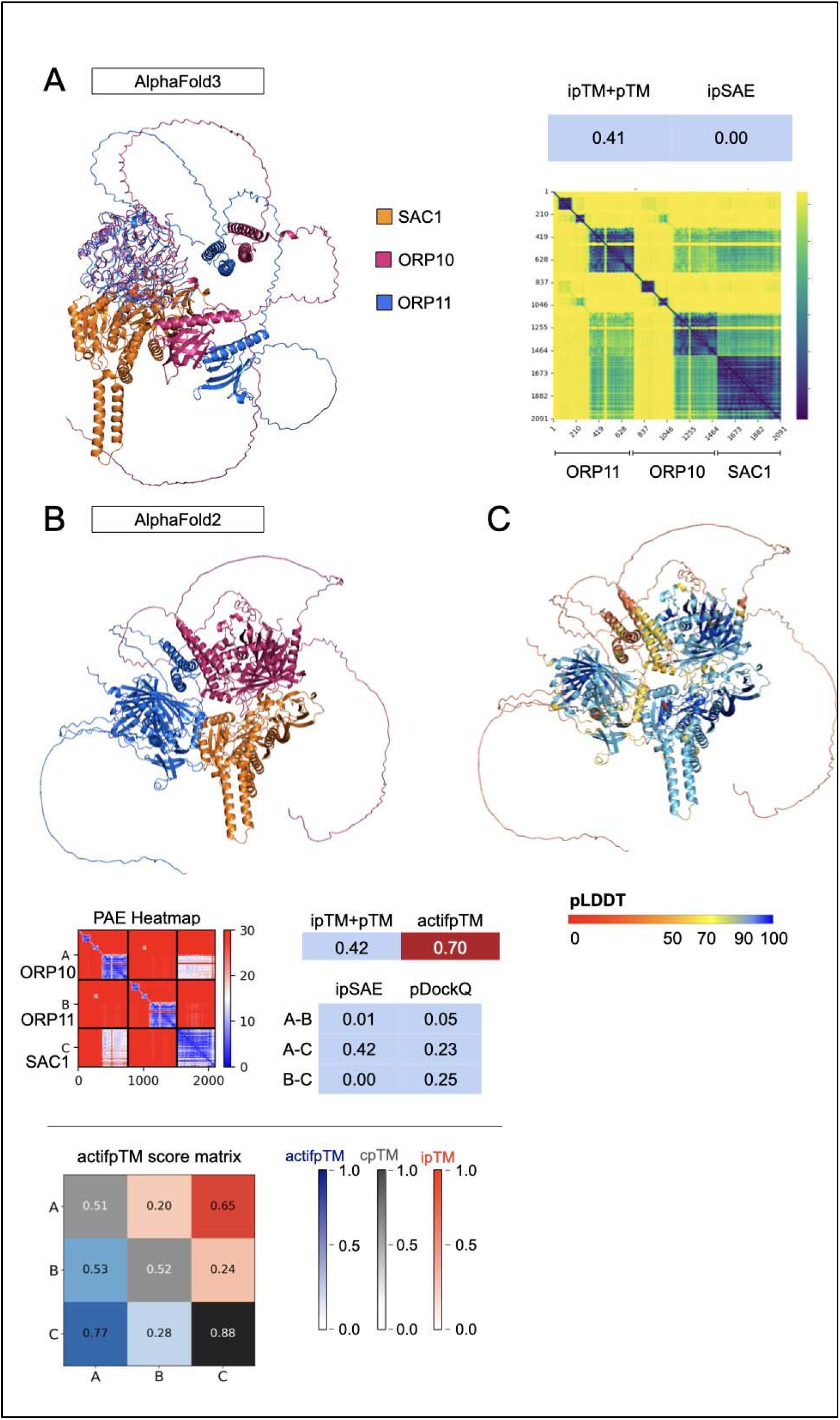
**A**. AlphaFold3 model of ORP10-ORP11-SAC1 coloured by chain. On the right, scoring metrics and PAE heatmap. **B**. AlphaFold2 model of ORP10-ORP11-SAC1 coloured by chain. On the right, scoring metrics and PAE heatmap. “ipTM+pTM” represents the weighted confidence score, calculated as 0.8 × ipTM + 0.2 × pTM. “Interface ipSAE” represents the ipSAE_d0dom score. **C**. AlphaFold2 model of ORP10-ORP11-SAC1 coloured by per-residue pLDDT confidence scores. Colour scheme follows the AlphaFold standard (credit: Konstantin Korotkov).

**Fig. S9.**
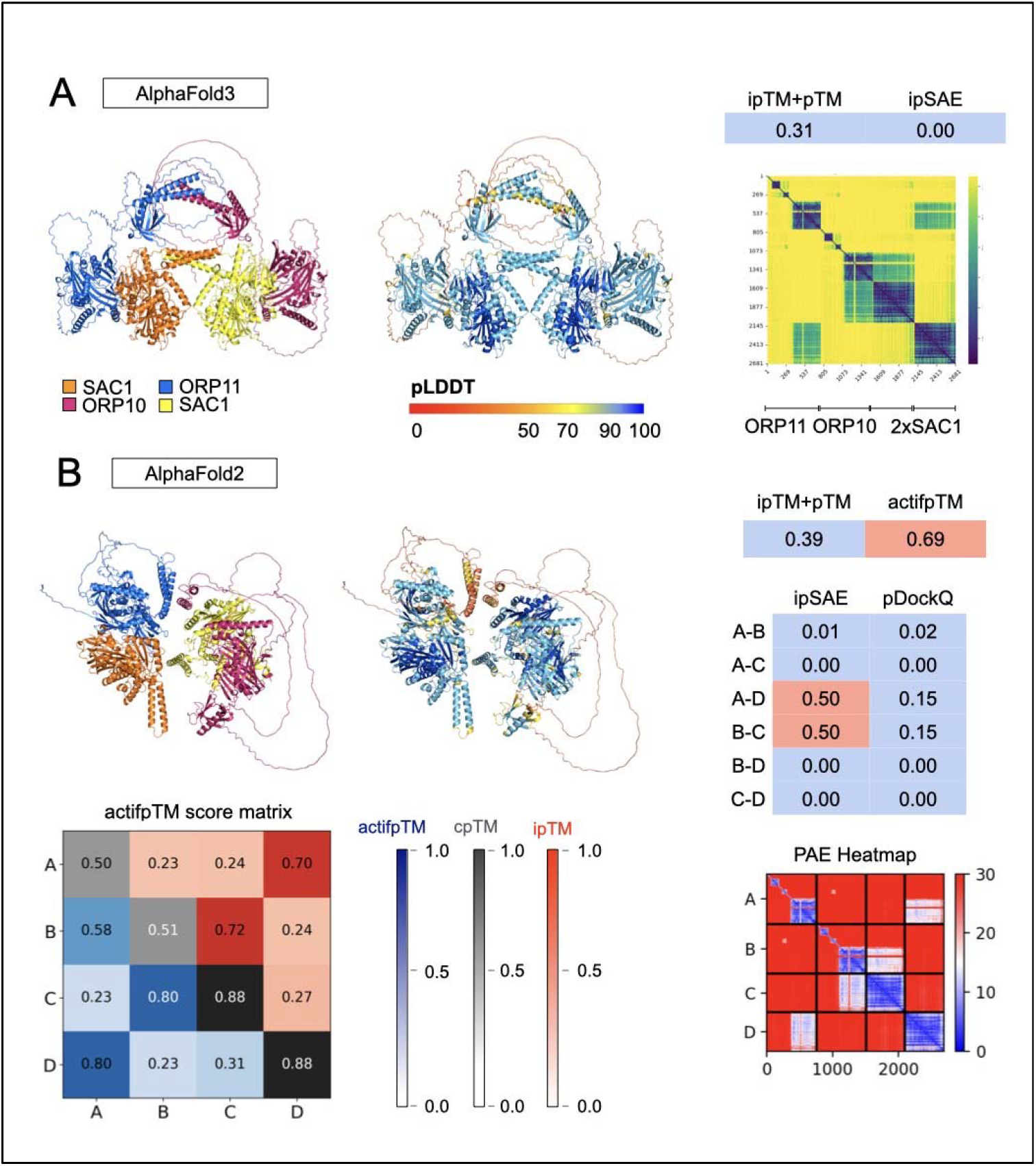
**A**. AlphaFold3 ORP10-ORP11 dimer modelled with two SAC1 proteins coloured by chain (left) and by per-residue pLDDT confidence scores (right). Colour scheme follows the AlphaFold standard (credit: Konstantin Korotkov). On the right, scoring metrics and PAE heatmap. **B**. AlphaFold2 ORP10-ORP11 dimer modelled with two SAC1 proteins coloured by chain (left) and by per-residue pLDDT confidence scores (right). On the right, scoring metrics and PAE heatmap. “ipTM+pTM” represents the weighted confidence score, calculated as 0.8 × ipTM + 0.2 × pTM. “Interface ipSAE” represents the ipSAE_d0dom score.

**Fig. S10.**
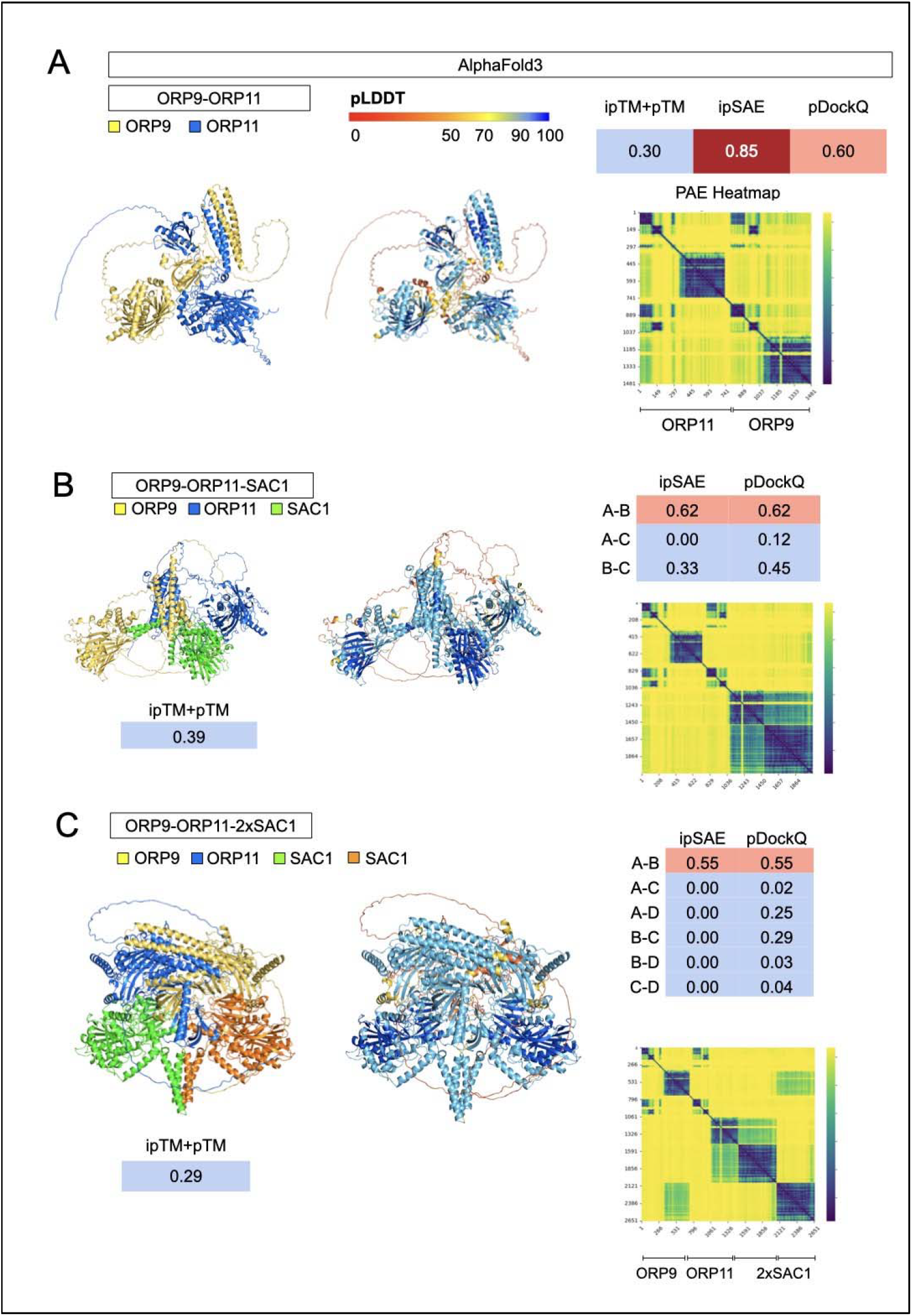
**A**. AlphaFold3 ORP9-ORP11 dimer, **B**. ORP9-ORP11 dimer with SAC1, and **C**. ORP9-ORP11 dimer modelled with two SAC1 proteins coloured by chain (left) and by per-residue pLDDT confidence scores (right). Colour scheme follows the AlphaFold standard (credit: Konstantin Korotkov). On the right, scoring metrics and PAE heatmap. “ipTM+pTM” represents the weighted confidence score, calculated as 0.8 × ipTM + 0.2 × pTM. “Interface ipSAE” represents the ipSAE_d0dom score.

**Fig. S11.**
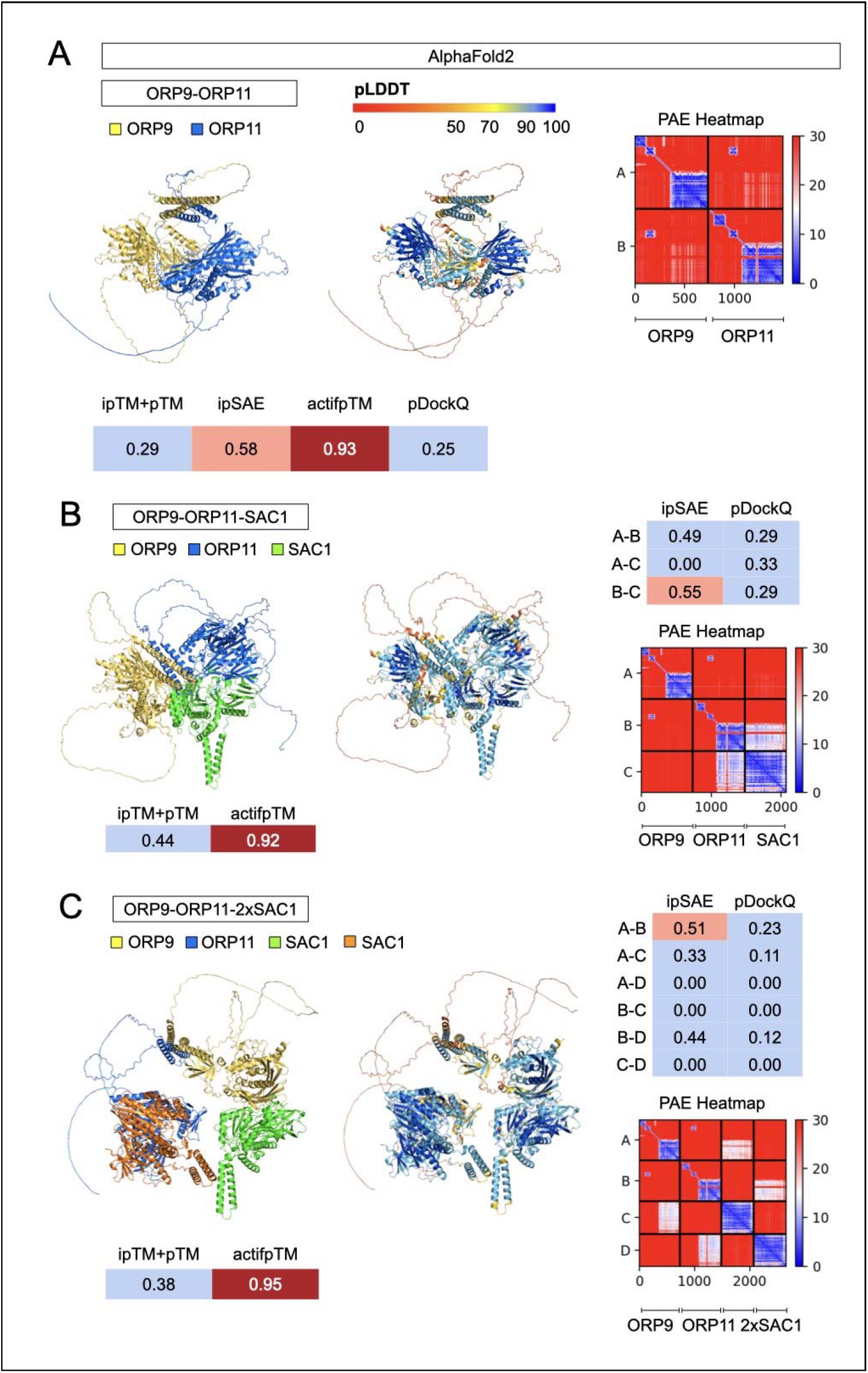
**A**. AlphaFold2-Multimer ORP9-ORP11 dimer, **B**. ORP9-ORP11 dimer with SAC1, and **C**. ORP9-ORP11 dimer modelled with two SAC1 proteins coloured by chain (left) and by per-residue pLDDT confidence scores (right). Colour scheme follows the AlphaFold standard (credit: Konstantin Korotkov). On the right, scoring metrics and PAE heatmap. “ipTM+pTM” represents the weighted confidence score, calculated as 0.8 × ipTM + 0.2 × pTM. “Interface ipSAE” represents the ipSAE_d0dom score.

**Fig. S12.**
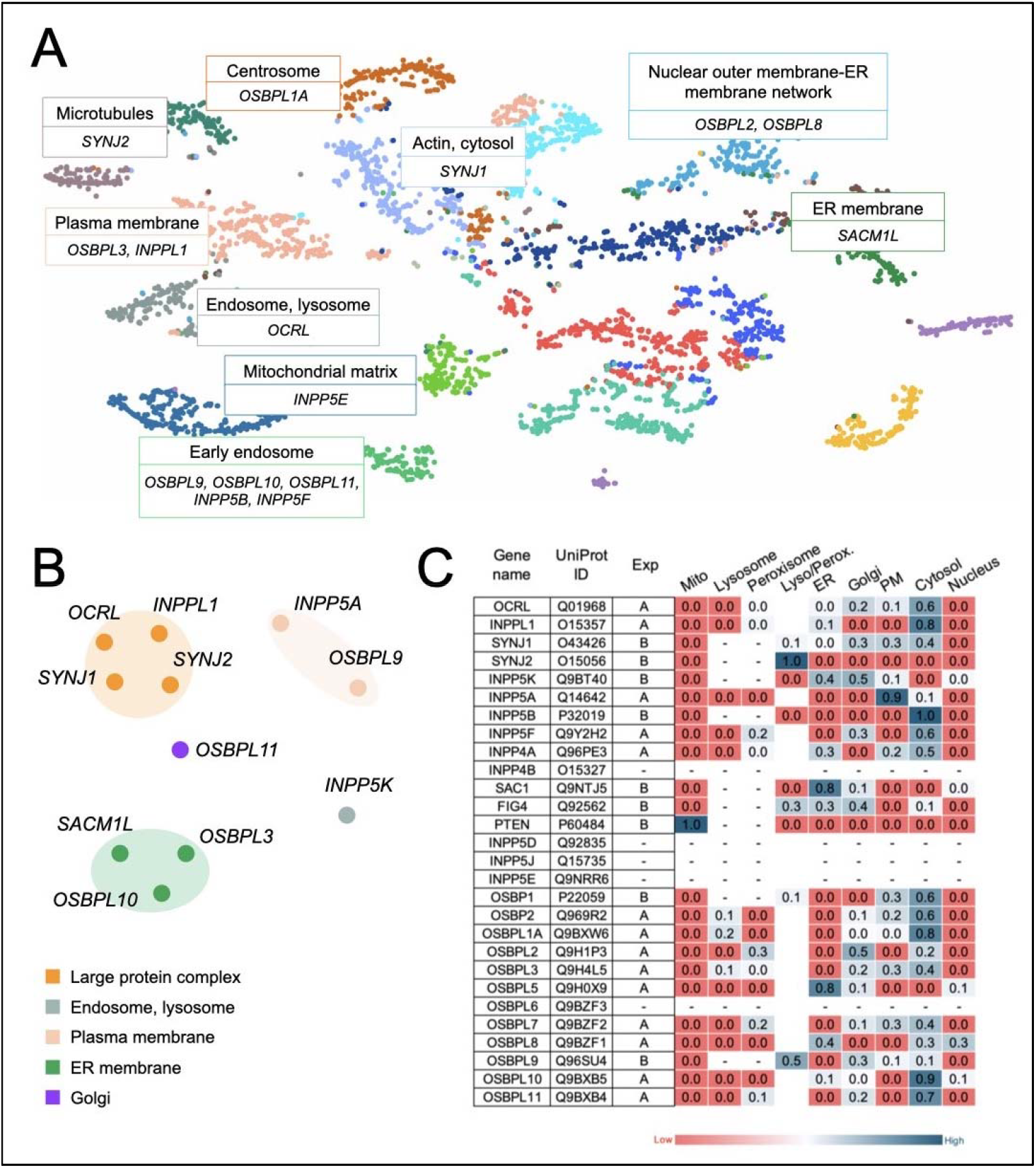
Subcellular localisation of ORP family proteins and PIPs. **A**. Human Cell Map dataset (adapted ^52^). **B**. Map of the Cell cellular compartment annotations in HeLa cells ^54^. **C**. Prolocate dataset (adapted ^53^), based on organelle fractionation of rat liver. Results are shown for two experimental conditions: A, fasted; B, fed.

## References

1. Prinz WA, Toulmay A, Balla T. The functional universe of membrane contact sites. Nat Rev Mol Cell Biol. Jan 2020;21(1):7–24. doi:10.1038/s41580-019-0180-9

2. Voeltz GK, Sawyer EM, Hajnoczky G, Prinz WA. Making the connection: How membrane contact sites have changed our view of organelle biology. Cell. Jan 18 2024;187(2):257–270. doi:10.1016/j.cell.2023.11.040

3. Saheki Y, De Camilli P. Endoplasmic Reticulum-Plasma Membrane Contact Sites. Annu Rev Biochem. Jun 20 2017;86:659–684. doi:10.1146/annurev-biochem061516-044932

4. Di Paolo G, De Camilli P. Phosphoinositides in cell regulation and membrane dynamics. Nature. Oct 12 2006;443(7112):651–7. doi:10.1038/nature05185

5. Posor Y, Jang W, Haucke V. Phosphoinositides as membrane organizers. Nat Rev Mol Cell Biol. Dec 2022;23(12):797–816. doi:10.1038/s41580-022-00490-x

6. Balla T. Phosphoinositides: tiny lipids with giant impact on cell regulation. Physiol Rev. Jul 2013;93(3):1019–137. doi:10.1152/physrev.00028.2012

7. Balla T, Gulyas G, Kim YJ, Pemberton J. Phosphoinositides and Calcium Signaling. A Marriage Arranged in Er-Pm Contact Sites. Curr Opin Physiol. Oct 2020;17:149–157. doi:10.1016/j.cophys.2020.08.007

8. Davies EM, Jones EI, Ooms LM, Gurung R, McGrath MJ, Mitchell CA. Phosphoinositide phosphatases: Modifiers of phosphoinositide signaling in health and disease. Biochim Biophys Acta Mol Cell Biol Lipids. Aug 2025;1870(6):159652. doi:10.1016/j.bbalip.2025.159652

9. Faucherre A, Desbois P, Satre V, Lunardi J, Dorseuil O, Gacon G. Lowe syndrome protein OCRL1 interacts with Rac GTPase in the trans-Golgi network. Hum Mol Genet. Oct 1 2003;12(19):2449–56. doi:10.1093/hmg/ddg250

10. Dong R, Zhu T, Benedetti L, et al. The inositol 5-phosphatase INPP5K participates in the fine control of ER organization. J Cell Biol. Oct 1 2018;217(10):3577–3592. doi:10.1083/jcb.201802125

11. Yang Y, Wang G, Huang X, Du Z. Crystallographic and modelling studies suggest that the SKICH domains from different protein families share a common Ig-like fold but harbour substantial structural variations. J Biomol Struct Dyn. 2015;33(7):1385–98. doi:10.1080/07391102.2014.951688

12. Zewe JP, Wills RC, Sangappa S, Goulden BD, Hammond GR. SAC1 degrades its lipid substrate PtdIns4P in the endoplasmic reticulum to maintain a steep chemical gradient with donor membranes. Elife. Feb 20 2018;7 doi:10.7554/eLife.35588

13. Cabukusta B, Borst Pauwels S, Akkermans J, et al. The ORP9-ORP11 dimer promotes sphingomyelin synthesis. Elife. Aug 6 2024;12 doi:10.7554/eLife.91345

14. Kawasaki A, Sakai A, Nakanishi H, et al. PI4P/PS countertransport by ORP10 at ER-endosome membrane contact sites regulates endosome fission. J Cell Biol. Jan 3 2022;221(1) doi:10.1083/jcb.202103141

15. Nakatsu F, Kawasaki A. Functions of Oxysterol-Binding Proteins at Membrane Contact Sites and Their Control by Phosphoinositide Metabolism. Front Cell Dev Biol. 2021;9:664788. doi:10.3389/fcell.2021.664788

16. Olkkonen VM, Li S. Oxysterol-binding proteins: sterol and phosphoinositide sensors coordinating transport, signaling and metabolism. Prog Lipid Res. Oct 2013;52(4):529–38. doi:10.1016/j.plipres.2013.06.004

17. Zhou Y, Li S, Mayranpaa MI, et al. OSBP-related protein 11 (ORP11) dimerizes with ORP9 and localizes at the Golgi-late endosome interface. Exp Cell Res. Nov 15 2010;316(19):3304–16. doi:10.1016/j.yexcr.2010.06.008

18. Murphy SE, Levine TP. VAP, a Versatile Access Point for the Endoplasmic Reticulum: Review and analysis of FFAT-like motifs in the VAPome. Biochim Biophys Acta. Aug 2016;1861(8 Pt B):952–961. doi:10.1016/j.bbalip.2016.02.009

19. Du X, Kumar J, Ferguson C, et al. A role for oxysterol-binding protein-related protein 5 in endosomal cholesterol trafficking. J Cell Biol. Jan 10 2011;192(1):121–35. doi:10.1083/jcb.201004142

20. Sohn M, Korzeniowski M, Zewe JP, et al. PI(4,5)P(2) controls plasma membrane PI4P and PS levels via ORP5/8 recruitment to ER-PM contact sites. J Cell Biol. May 7 2018;217(5):1797–1813. doi:10.1083/jcb.201710095

21. Del Bel LM, Brill JA. Sac1, a lipid phosphatase at the interface of vesicular and nonvesicular transport. Traffic. May 2018;19(5):301–318. doi:10.1111/tra.12554

22. Moser von Filseck J, Vanni S, Mesmin B, Antonny B, Drin G. A phosphatidylinositol-4-phosphate powered exchange mechanism to create a lipid gradient between membranes. Nat Commun. Apr 7 2015;6:6671. doi:10.1038/ncomms7671

23. Manford A, Xia T, Saxena AK, et al. Crystal structure of the yeast Sac1: implications for its phosphoinositide phosphatase function. EMBO J. May 5 2010;29(9):1489–98. doi:10.1038/emboj.2010.57

24. Schafer JH, Korner C, Esch BM, et al. Structure of the ceramide-bound SPOTS complex. Nat Commun. Oct 4 2023;14(1):6196. doi:10.1038/s41467-023-41747-z

25. Stefan CJ, Manford AG, Baird D, Yamada-Hanff J, Mao Y, Emr SD. Osh proteins regulate phosphoinositide metabolism at ER-plasma membrane contact sites. Cell. Feb 4 2011;144(3):389–401. doi:10.1016/j.cell.2010.12.034

26. Molodenskiy D, Maurer VJ, Yu D, et al. AlphaPulldown2-a general pipeline for high-throughput structural modeling. Bioinformatics. Mar 4 2025;41(3) doi:10.1093/bioinformatics/btaf115

27. Dunbrack RL, Jr. Res ipSAE loquunt: What’s wrong with AlphaFold’s ipTM score and how to fix it. bioRxiv. Feb 14 2025; doi:10.1101/2025.02.10.637595

28. Varga JK, Ovchinnikov S, Schueler-Furman O. actifpTM: a refined confidence metric of AlphaFold2 predictions involving flexible regions. Bioinformatics. Mar 4 2025;41(3) doi:10.1093/bioinformatics/btaf107

29. Jumper J, Evans R, Pritzel A, et al. Highly accurate protein structure prediction with AlphaFold. Nature. Aug 2021;596(7873):583–589. doi:10.1038/s41586-021-03819-2

30. Mirdita M, Schutze K, Moriwaki Y, Heo L, Ovchinnikov S, Steinegger M. ColabFold: making protein folding accessible to all. Nat Methods. Jun 2022;19(6):679–682. doi:10.1038/s41592-022-01488-1

31. Bret H, Gao J, Zea DJ, Andreani J, Guerois R. From interaction networks to interfaces, scanning intrinsically disordered regions using AlphaFold2. Nat Commun. Jan 18 2024;15(1):597. doi:10.1038/s41467-023-44288-7

32. Lee CY, Hubrich D, Varga JK, et al. Systematic discovery of protein interaction interfaces using AlphaFold and experimental validation. Mol Syst Biol. Feb 2024;20(2):75–97. doi:10.1038/s44320-023-00005-6

33. Teufel F, Refsgaard JC, Kasimova MA, et al. Deorphanizing Peptides Using Structure Prediction. J Chem Inf Model. May 8 2023;63(9):2651–2655. doi:10.1021/acs.jcim.3c00378

34. Abramson J, Adler J, Dunger J, et al. Accurate structure prediction of biomolecular interactions with AlphaFold 3. Nature. Jun 2024;630(8016):493–500. doi:10.1038/s41586-024-07487-w

35. Kim W, Mirdita M, Levy Karin E, et al. Rapid and sensitive protein complex alignment with Foldseek-Multimer. Nat Methods. Mar 2025;22(3):469–472. doi:10.1038/s41592-025-02593-7

36. Bryant P, Pozzati G, Elofsson A. Improved prediction of protein-protein interactions using AlphaFold2. Nat Commun. Mar 10 2022;13(1):1265. doi:10.1038/s41467-022-28865-w

37. Basu S, Wallner B. DockQ: A Quality Measure for Protein-Protein Docking Models. PLoS One. 2016;11(8):e0161879. doi:10.1371/journal.pone.0161879

38. Ahmad A. AF3 Results Visualization. Accessed 1 Nov 2024, https://colab.research.google.com/github/Ash100/Biopython/blob/main/AF3_Results_Visualization.ipynb#scrollTo=YZAVOqx6dig5

39. Krissinel E. Crystal contacts as nature’s docking solutions. J Comput Chem. Jan 15 2010;31(1):133–43. doi:10.1002/jcc.21303

40. Krissinel E, Henrick K. Inference of macromolecular assemblies from crystalline state. J Mol Biol. Sep 21 2007;372(3):774–97. doi:10.1016/j.jmb.2007.05.022

41. Álvarez-Salmoral D, Borza R, Xie R, Joosten RP, Hekkelman ML, Perrakis A. AlphaBridge: tools for the analysis of predicted macromolecular complexes. Preprint. bioRxiv. 2024; doi:10.1101/2024.10.23.619601

42. Zhu W, Shenoy A, Kundrotas P, Elofsson A. Evaluation of AlphaFold-Multimer prediction on multi-chain protein complexes. Bioinformatics. Jul 1 2023;39(7) doi:10.1093/bioinformatics/btad424

43. Barrio-Hernandez I, Yeo J, Janes J, et al. Clustering predicted structures at the scale of the known protein universe. Nature. Oct 2023;622(7983):637–645. doi:10.1038/s41586-023-06510-w

44. van Kempen M, Kim SS, Tumescheit C, et al. Fast and accurate protein structure search with Foldseek. Nat Biotechnol. Feb 2024;42(2):243–246. doi:10.1038/s41587-023-01773-0

45. Jo S, Kim T, Im W. Automated builder and database of protein/membrane complexes for molecular dynamics simulations. PLoS One. Sep 12 2007;2(9):e880. doi:10.1371/journal.pone.0000880

46. Jo S, Kim T, Iyer VG, Im W. CHARMM-GUI: a web-based graphical user interface for CHARMM. J Comput Chem. Aug 2008;29(11):1859–65. doi:10.1002/jcc.20945

47. Wu EL, Cheng X, Jo S, et al. CHARMM-GUI Membrane Builder toward realistic biological membrane simulations. J Comput Chem. Oct 15 2014;35(27):1997–2004. doi:10.1002/jcc.23702

48. Lomize MA, Pogozheva ID, Joo H, Mosberg HI, Lomize AL. OPM database and PPM web server: resources for positioning of proteins in membranes. Nucleic Acids Res. Jan 2012;40(Database issue):D370–6. doi:10.1093/nar/gkr703

49. Lee J, Cheng X, Swails JM, et al. CHARMM-GUI Input Generator for NAMD, GROMACS, AMBER, OpenMM, and CHARMM/OpenMM Simulations Using the CHARMM36 Additive Force Field. J Chem Theory Comput. Jan 12 2016;12(1):405–13. doi:10.1021/acs.jctc.5b00935

50. Landau M, Mayrose I, Rosenberg Y, et al. ConSurf 2005: the projection of evolutionary conservation scores of residues on protein structures. Nucleic Acids Res. Jul 1 2005;33(Web Server issue):W299–302. doi:10.1093/nar/gki370

51. Krapp LF, Abriata LA, Cortes Rodriguez F, Dal Peraro M. PeSTo: parameter-free geometric deep learning for accurate prediction of protein binding interfaces. Nat Commun. Apr 18 2023;14(1):2175. doi:10.1038/s41467-023-37701-8

52. Go CD, Knight JDR, Rajasekharan A, et al. A proximity-dependent biotinylation map of a human cell. Nature. Jul 2021;595(7865):120–124. doi:10.1038/s41586-021-03592-2

53. Jadot M, Boonen M, Thirion J, et al. Accounting for Protein Subcellular Localization: A Compartmental Map of the Rat Liver Proteome. Mol Cell Proteomics. Feb 2017;16(2):194–212. doi:10.1074/mcp.M116.064527

54. Itzhak DN, Tyanova S, Cox J, Borner GH. Global, quantitative and dynamic mapping of protein subcellular localization. Elife. Jun 9 2016;5 doi:10.7554/eLife.16950

55. Pereira GP, Gouzien C, Souza PCT, Martin J. Challenges in predicting PROTAC-mediated protein-protein interfaces with AlphaFold reveal a general limitation on small interfaces. Bioinform Adv. 2025;5(1):vbaf056. doi:10.1093/bioadv/vbaf056

56. Bastian M, Heymann S, Jacomy M. Gephi: An Open Source Software for Exploring and Manipulating Networks. Conference paper. International AAAI Conference on Weblogs and Social Media. 2009; doi:10.1609/icwsm.v3i1.13937

57. Nam KH. Evaluation of AlphaFold3 for the fatty acids docking to human fatty acid-binding proteins. J Mol Graph Model. Dec 2024;133:108872. doi:10.1016/j.jmgm.2024.108872

58. Gautier R, Douguet D, Antonny B, Drin G. HELIQUEST: a web server to screen sequences with specific alpha-helical properties. Bioinformatics. Sep 15 2008;24(18):2101–2. doi:10.1093/bioinformatics/btn392

59. Kara B, Koroglu C, Peltonen K, et al. Severe neurodegenerative disease in brothers with homozygous mutation in POLR1A. Eur J Hum Genet. Feb 2017;25(3):315–323. doi:10.1038/ejhg.2016.183

60. Yang Z, Zeng X, Zhao Y, Chen R. AlphaFold2 and its applications in the fields of biology and medicine. Signal Transduct Target Ther. Mar 14 2023;8(1):115. doi:10.1038/s41392-023-01381-z

61. Abdelkareem AO, Hall CF, Marutharaj A, Narta K, Verhey TB, Morrissy AS. XenoSignal: Investigating Intra- and Inter-Species Ligand-Receptor Interactions Using AlphaFold3. bioRxiv. 2025; doi:10.1101/2025.08.13.670200

62. Catoiu EA, Kambo D, Rodriguez B, et al. QSProteome: A Community-Driven Interactive Platform for Large-Scale Exploration and Evaluation of Predicted Protein Complex Structures. Pre-print. bioRxiv. 2025; doi:10.1101/2025.09.10.675416

63. Mathew VS, Kellogg GD, Lai WKM. Protein structure prediction and design for high-throughput computing. Pre-print. bioRxiv. 2025; doi:10.1101/2025.07.18.665594

64. Fernandez-Busnadiego R, Saheki Y, De Camilli P. Three-dimensional architecture of extended synaptotagmin-mediated endoplasmic reticulum-plasma membrane contact sites. Proc Natl Acad Sci U S A. Apr 21 2015;112(16):E2004–13. doi:10.1073/pnas.1503191112

65. Ghaemmaghami S, Huh WK, Bower K, et al. Global analysis of protein expression in yeast. Nature. Oct 16 2003;425(6959):737–41. doi:10.1038/nature02046

66. Knodler A, Konrad G, Mayinger P. Expression of yeast lipid phosphatase Sac1p is regulated by phosphatidylinositol-4-phosphate. BMC Mol Biol. Jan 28 2008;9:16. doi:10.1186/1471-2199-9-16

67. Huang X, Jiang C, Yu L, Yang A. Current and Emerging Approaches for Studying Inter-Organelle Membrane Contact Sites. Front Cell Dev Biol. 2020;8:195. doi:10.3389/fcell.2020.00195

68. Jing J, Liu G, Huang Y, Zhou Y. A molecular toolbox for interrogation of membrane contact sites. J Physiol. May 2020;598(9):1725–1739. doi:10.1113/JP277761

69. Mondal S, Narayan KB, Powers I, Botterbusch S, Baumgart T. Endophilin recruitment drives membrane curvature generation through coincidence detection of GPCR loop interactions and negative lipid charge. J Biol Chem. Jan-Jun 2021;296:100140. doi:10.1074/jbc.RA120.016118

70. Vicinanza M, Puri C, Rubinsztein DC. Coincidence detection of RAB11A and PI(3)P by WIPI2 directs autophagosome formation. Oncotarget. Apr 5 2019;10(27):2579–2580. doi:10.18632/oncotarget.26829

71. Chou JC, Chatterjee P, Decosto CM, Dassama LMK. A machine learning model for the proteome-wide prediction of lipid-interacting proteins. bioRxiv. May 25 2025; doi:10.1101/2024.01.26.577452

72. He R, Liu F, Wang H, et al. ORP9 and ORP10 form a heterocomplex to transfer phosphatidylinositol 4-phosphate at ER-TGN contact sites. Cell Mol Life Sci. Feb 28 2023;80(3):77. doi:10.1007/s00018-023-04728-5

73. Tan JX, Finkel T. A phosphoinositide signalling pathway mediates rapid lysosomal repair. Nature. Sep 2022;609(7928):815–821. doi:10.1038/s41586-022-05164-4

74. Tauriello G, Waterhouse AM, Haas J, et al. ModelArchive: A Deposition Database for Computational Macromolecular Structural Models. J Mol Biol. Aug 1 2025;437(15):168996. doi:10.1016/j.jmb.2025.168996

